# An anchored experimental design and meta-analysis approach to address batch effects in large-scale metabolomics

**DOI:** 10.1101/2022.03.25.485859

**Authors:** Amanda O. Shaver, Brianna M. Garcia, Goncalo J. Gouveia, Alison M. Morse, Zihao Liu, Carter K. Asef, Ricardo M. Borges, Franklin E. Leach, Erik C. Andersen, I. Jonathan Amster, Facundo M. Fernández, Arthur S. Edison, Lauren M. McIntyre

**Author notes:** **Corresponding author:** Lauren M. McIntyre, Department of Molecular Genetics and Microbiology, University of Florida Genetics Institute, University of Florida, Mowry Road. These authors contributed equally (Amanda O. Shaver, Brianna M. Garcia).

## Abstract

Large-scale untargeted metabolomics studies suffer from individual variation, batch effects and instrument variability, making comparisons of common spectral features across studies difficult. One solution is to compare studies after compound identification. However, compound identification is expensive and time consuming. We successfully identify common spectral features across multiple studies, with a generalizable experimental design approach. First, we included an anchor strain, PD1074, during sample and data collection. Second, we collected data in blocks with multiple controls. These anchors enabled us to successfully integrate three studies of *Caenorhabditis elegans* for nuclear magnetic resonance (NMR) spectroscopy and liquid chromatography-mass spectrometry (LC-MS) data from five different assays. We found 34% and 14% of features to be significant in LC-MS and NMR, respectively. Between 20-50% of spectral features differ in a mutant and among a set of genetically diverse natural strains, suggesting this reduced set of spectral features are excellent targets for compound identification.

**GRAPHICAL ABSTRACT:** 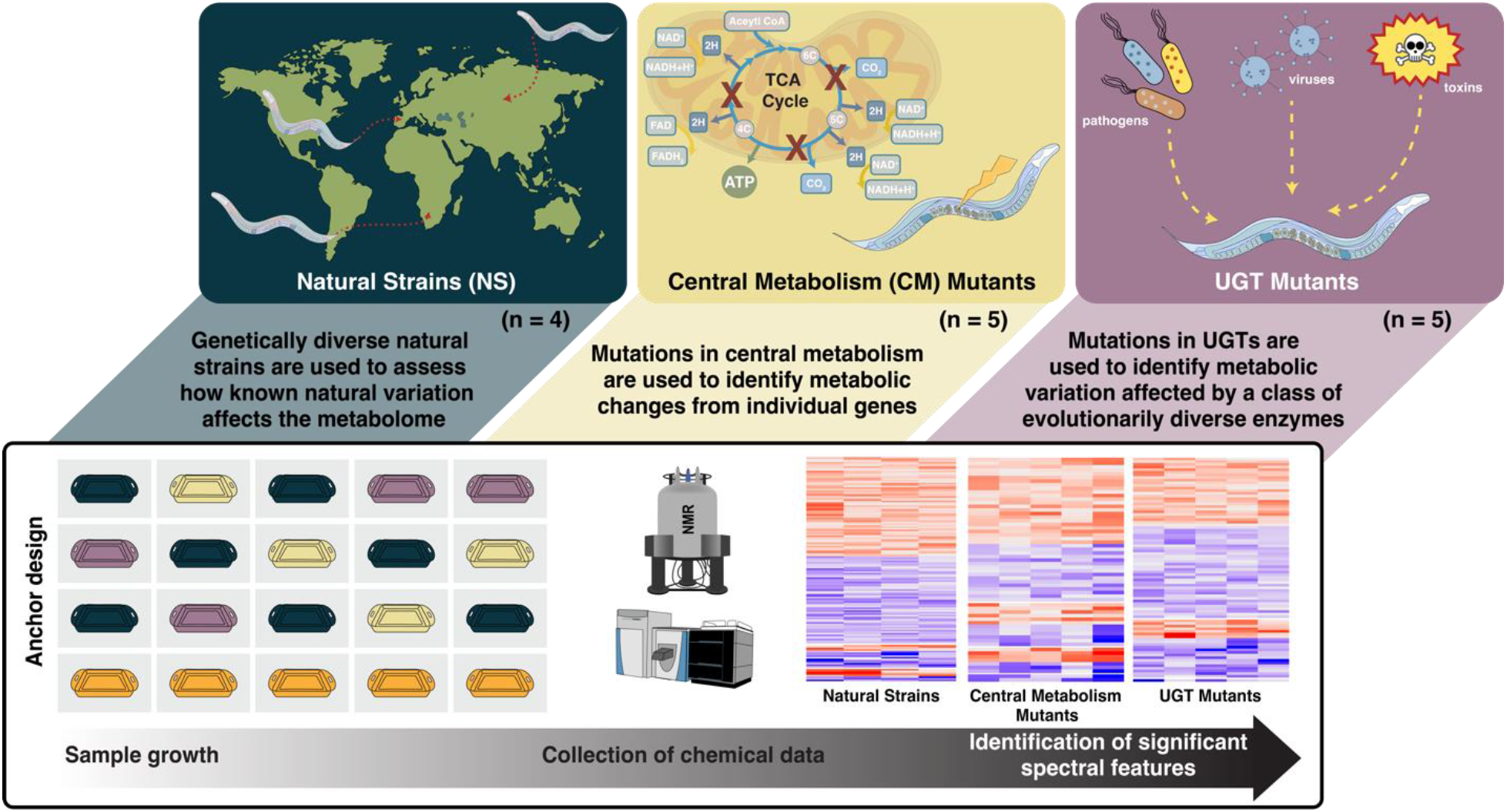

Fourteen *C. elegans* strains are used in three individual studies. PD1074, the anchor control strain (orange), is grown alongside test strains (green, yellow, purple). Multiple biological replicates of PD1074 captures environmental variation in growth conditions. Non-polar and polar metabolic data across the three studies (*i*.*e*., natural strains, central metabolism mutants, and UGT mutants) were collected by nuclear magnetic resonance (NMR) spectroscopy and liquid chromatography-mass spectrometry (LC-MS). Data acquisition controls in each block included biological reference material and pooled PD1074 samples. Biological replicates of PD1074 (n = 42 for LC-MS, n = 52 for NMR) were included in all batches. Meta-analysis provided comparable inferences to mixed effects models, and the estimated relative effects of each test strain to PD1074 and straightforward comparisons of test strains across experiments.

---

Untargeted metabolomics studies compare the variation in small molecules caused by genetic perturbations, treatments, and environmental differences^1^. Metabolomics is a powerful tool in biomarker discovery and holds great promise for precision medicine^2-4^. Targeted metabolomics is common in studies exploring human health questions that range from aging^5, 6^ to complex diseases^7-12^. An advantage of untargeted metabolomics for these questions is the ability to reach beyond sets of well studied compounds to explore differences in an unbiased way^13^. Despite the attractiveness of an unbiased survey, untargeted metabolomics has well known challenges. In particular, the collection of highly variable biological material in a reproducible manner across batches makes the identification of differential compounds and comparisons of their abundances across datasets challenging. Chemical annotation of compounds, which is key to combining data across studies, requires considerable time and labor^14^. Given this bottleneck, it is essential to find novel ways to prioritize spectral features and overcome intractable challenges such as matrix effects, instrument drift, and batch variation^15-18^.

Batch effects across experiments are an enormous problem in untargeted metabolomics and a barrier to adopting these methods^19^. Normalizing to a quality control (QC) or biological reference material (BRM) material included in each batch has been shown to be effective^10, 15, 16^. Although normalization strategies are improving^15, 16^; non-linear effects^20^, sample variation, the inability to separate environmental variance, and analytical artifacts^17^ still pose ongoing challenges to the identification of common spectral features across studies. While different approaches to sample-based and data-based normalization have been described, such as total protein content, total ion count (TIC), and pooled QCs^12, 21^, reproducibility and heteroscedasticity (unequal variance) issues remain problematic^22-26^.

Our goal, common to many studies, is to compare groups across large numbers of independent samples^27-30^. As sample size increases, challenges associated with variation must be accounted for appropriately. In metabolomics studies, variation in pre-analytical sample collection (growth), analytical sample preparation (extraction), and data collection (instrument)^31^ can be confounded (**Figure 1**). Identification of shared spectral features using a BRM is a successful strategy^31, 32^ that has proven essential in large-scale studies^32-34^. Implementation of BRM controls for instrument variation can estimate and normalize extraction variation^16, 18, 31^. However, variation among samples within the group remains. Metabolites may only be present in some samples or some batches. In both liquid chromatography-mass spectrometry (LC-MS) and nuclear magnetic resonance (NMR) spectroscopy, ambiguity in whether features are generated by genetic or environmental factors coupled with batch effects and challenges in peak picking algorithms present obstacles to apply untargeted metabolomics to broader studies^17, 35^.

**Figure 1.**
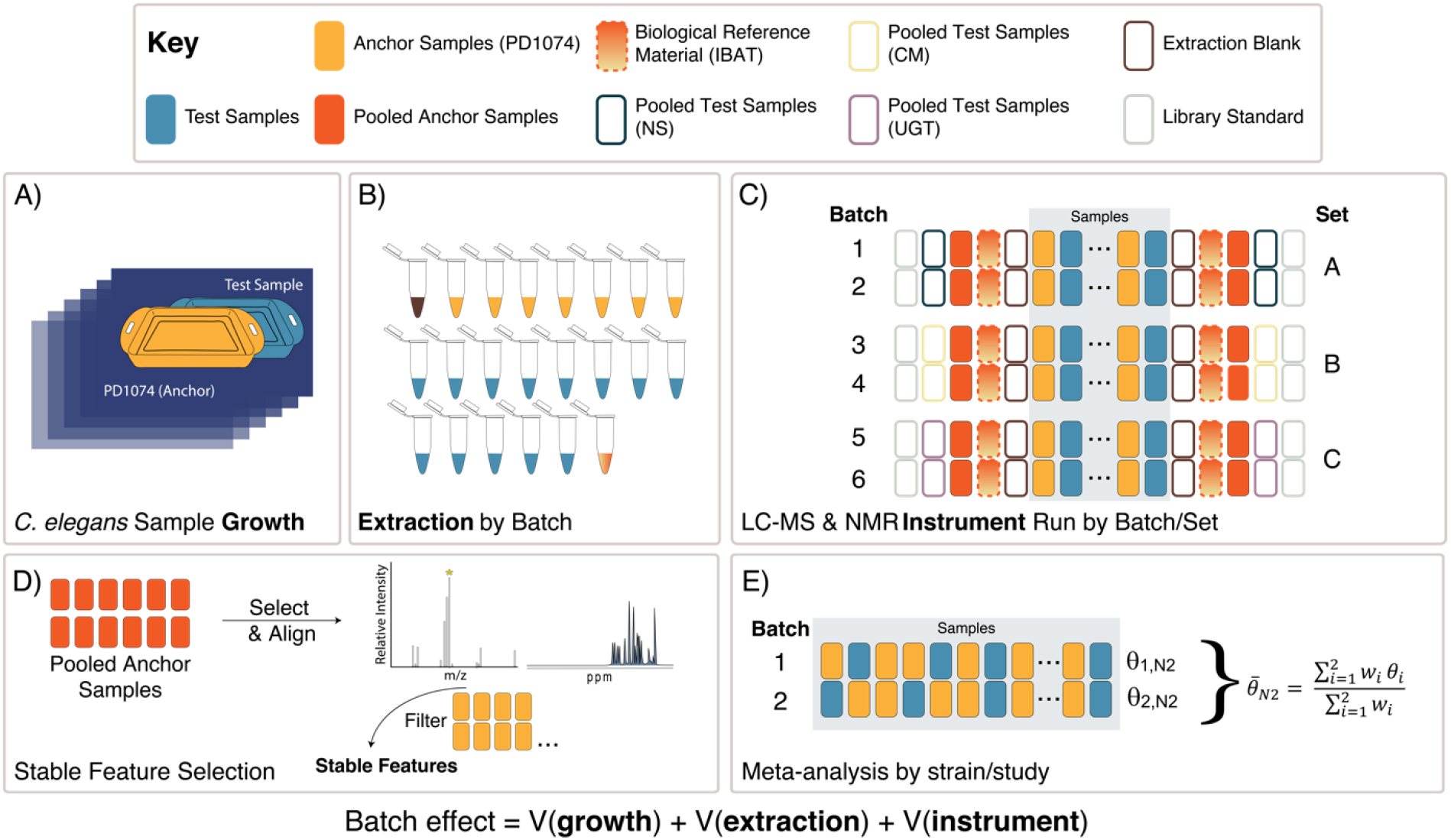
Experimental design overview. **(A)** Each *C. elegans* LSCP was grown and harvested with at least one PD1074 sample (sample growth variation captured) (**Figure S2**). **(B)** Multiple independent PD1074 samples and test strains (NS, CM mutants, or UGT mutants), IBAT references, and blanks were included in each batch for LC-MS or NMR (batch preparation variation captured). **(C)** A total of six batches in three sets were collected. Instrument controls, library standards, and replicate measurements of the pooled PD1074 samples (instrumentation variation captured) were in each run. Each test strain was collected in two independent sequential batches and a pool of all test samples are measured multiple times. **(D)** In LC-MS, PD1074 spectral features were first identified from PD1074 pools and retained if present above the level of the blank in 100% of the individual PD1074 spectra. In NMR, semi-automated peak-picking and binning were performed to extract peak heights and identify stable peaks present in PD1074 samples. **(E**) Data analysis was performed using meta-analysis models to identify spectral features of interest.

Although tools to handle extraction and instrumentation variation exist, their utility in large studies for samples with complex matrices is limited^33, 35, 36^. Here, we use the model system *Caenorhabditis elegans* to demonstrate that an augmented design combined with experimental blocks^37-39^ can be used to anchor studies and enable comparisons of stable spectral features across time without the need for compound identification.

*C. elegans* is a model organism ideally suited to study conserved small molecules in metabolism^40-42^. The worm’s short life cycle, self-fertilization of homozygous hermaphroditic individuals, ease of cultivation, and ability to propagate large numbers of animals^43^ are ideal for large-scale studies^42, 44-46^. These traits allow one to (i) develop, test, and validate approaches to identify stable spectral features, (ii) demonstrate the feasibility of large-scale biochemical pathway analyses with genetic mutants, and (iii) focus on spectral features likely to reveal essential components of metabolic pathways by comparing features that vary due to genetic perturbations.

We designed three *C. elegans* studies to link natural and deliberate knock-out genetic perturbations. The first and second studies comprised central metabolism (CM) mutants and UDP-glycosyltransferases (UGT) mutants as examples of primary and secondary metabolism, respectively. CM mutants have been used in studies showing that diagnostic changes can be associated with human disease^47, 48^. UGTs are an evolutionarily diverse class of Phase 2 enzymes involved in detoxification^49, 50^. Although UGTs are vital to internal detoxification across species, the functions of UGTs have not been well described ^49-52^. The third study comprises genetically diverse natural strains (NS) from a broad geographic base, used to describe natural variation in the metabolome of *C*. elegans^53^ including N2, a widely used laboratory-adapted strain^54^.

Collectively, CM and UGT mutants, and NS, allow us to (i) identify spectral features that vary due to genetic perturbations, (ii) compare the same spectral features across all three studies without compound identification, and (iii) plan future experiments that can be directly compared to these studies. The experimental design used here is straightforward to execute in model systems. One (or more) anchor strains (PD1074 here) are included alongside every test strain during growth and data collection, augmenting the design. Including the same strain enables measurement of the variation due to non-genetic effects. Augmented designs are common in large scale agricultural studies and they are used to compare large numbers of genotypes across heterogeneous environments^38, 39^. The inclusion of multiple biological replicates of the same strain during data acquisition enables the identification of stable features across a wide range of environmental conditions. Given the limited resources and expense of compound identification, analysis of the set of stable spectral features for differences in intensity in several contexts provides one way of prioritizing interesting compounds for identification.

## RESULTS

Here, we provide a method to identify stable spectral features and identify differences between groups using a straightforward meta-analytic approach. This demonstration is comprised of 104 independent samples collected in three studies of two batches each to produce five analytical datasets (3 LC-MS and 2 NMR) from two complementary technologies commonly used in untargeted metabolomics (**Figures 1 and S1**).

Our first study comprised of CM mutants (n = 5) identifies spectral features involved in central metabolism. The second study, UGT mutants (n = 5), identifies spectral features affected by Phase 2 enzymes involved in the detoxification system. The third study, NS (n = 4), assesses natural genetic variation. Collectively, these studies represent common hypotheses of general interest to metabolomics and genetic researchers (See **Table S1** for full strain details).

Stringent quality assurance/quality controls (QA/QC) combined with a focus on spectral features consistently detected in PD1074 identified: 3953 spectral features in reverse phase (RP) LC-MS positive, 377 in RP LC-MS negative, 199 in hydrophilic interaction liquid chromatography (HILIC) LC-MS positive, 585 in NMR polar, and 487 in NMR non-polar. An instrument failure occurred during the collection of the HILIC negative data (see Methods).

LC-MS spectral features often vary across biological replicates. Additional complexities include retention time drift, batch effects, and algorithmic limitations in estimating peak abundances in complex spectra^26, 55-58^. Including multiple independent PD1074 samples and pooled PD1074 samples in each batch can mitigate these issues. We continuously seeded and harvested PD1074 every time test samples were seeded or harvested during the large-scale culture plate (LSCP) growth process^43^. These PD1074 samples anchor the three studies and enable inter-study comparisons^37, 39^.

### PD1074 samples are pooled to control for within batch instrument variability

PD1074 LSCPs are genetically identical, leading to the expectation that the spectral features present in each biological replicate are a result of the strain’s genetic composition as they are stable across an extensive range of growth conditions (samples collected over six months) (**Figure S2**). All biological replicates of PD1074 are extracted separately. There were also technical replicates (repeat extractions of the same LSCP in different batches, n=20 for NMR, n =18 for LC-MS) that enabled an additional QC. During data acquisition, we included multiple biological replicates of PD1074 and a pool of the PD1074 samples included in each batch (**Figure 1C**). Comparing the PD1074 pools (measured twice n=12) across all batches (n=6) enables the identification of spectral features present across instrument runs over several months. Further, the selection of features present in all biological replicates of PD1074 ensures stable features across a range of environmental conditions (**Figure 1**). Iterative batch average method (IBAT) controls in the NMR study (from PD1074) combined with the biological replicates of PD1074 and the PD1074 pools enabled us to estimate the relative contribution of extraction (∼40%), growth (∼60%), and instrument variance, as expected in NMR, was negligible^31^. We are also able to directly compare the stable features detected using the BRM approach to PD1074 batch pools. We found that 97% of the features overlap.

### Meta-analysis identifies differences in spectral features between test and reference strains without the need for complex normalization

For each spectral feature, the difference in effect between the PD1074 individual LSCP (n=6-10) and each test strain (n=2-6) was estimated for each batch. We identified statistically significant spectral features by performing a meta-analysis across the two batches for each test strain^59^ eliminating the need to estimate and normalize/remove batch variance^37^ (**Figure 1E**). We compare a meta-analysis with a linear models analysis^60^ (**Figure S3**) and demonstrate that the final inferences are very similar, as predicted in larger studies that have compared individual analyses and meta-analytic approaches^60^. An advantage of the meta-analysis is the ability to apply this technique generally, even when there may be complex patterns of variance across batches such as those present in large cohort studies and/or due to technical variation (*e*.*g*., after an instrument interruption). Effect sizes can be used to compare test strains when data acquisition occurs independently across time. Effect sizes calculated in the meta-analysis are comparable to those calculated by a linear model analysis, demonstrating the successful implementation of meta-analysis when sample sizes are small (**Figure S3**).

We see a similar pattern across platforms for the percentage of significant features identified across the three studies, with the highest percentage found in the RP LC-MS (-) dataset (**Figure 2A**). The highest percentage of significant spectral features was 58% in the CM mutation study. In the individual strains, the CM mutant, VC1265 (*pyk-1*) had the largest overall effect across platforms and fractions, followed by RB2347 (*idh-2*). AUM2073 (*unc-119*) and KJ550 (*aco-1*) had the smallest overall effects (**Figures 2B, 3, and S4**). For the UGT mutants, VC2512 (*ugt-60*) had the largest overall effect, followed by RB2607 (*ugt-49*). RB2011 (*ugt-62*) had the smallest overall effect (**Figures 2C and S5**). These patterns demonstrate the variation in single knockouts of different genes. In the NS, the most genetically divergent strains from PD1074 (CB4856 and DL238) had the largest overall effect in both platforms, and N2 had a small set of differences, as expected, since PD1074 is a trackable variant of N2 (**Figures 2D and S6**).

**Figure 2.**
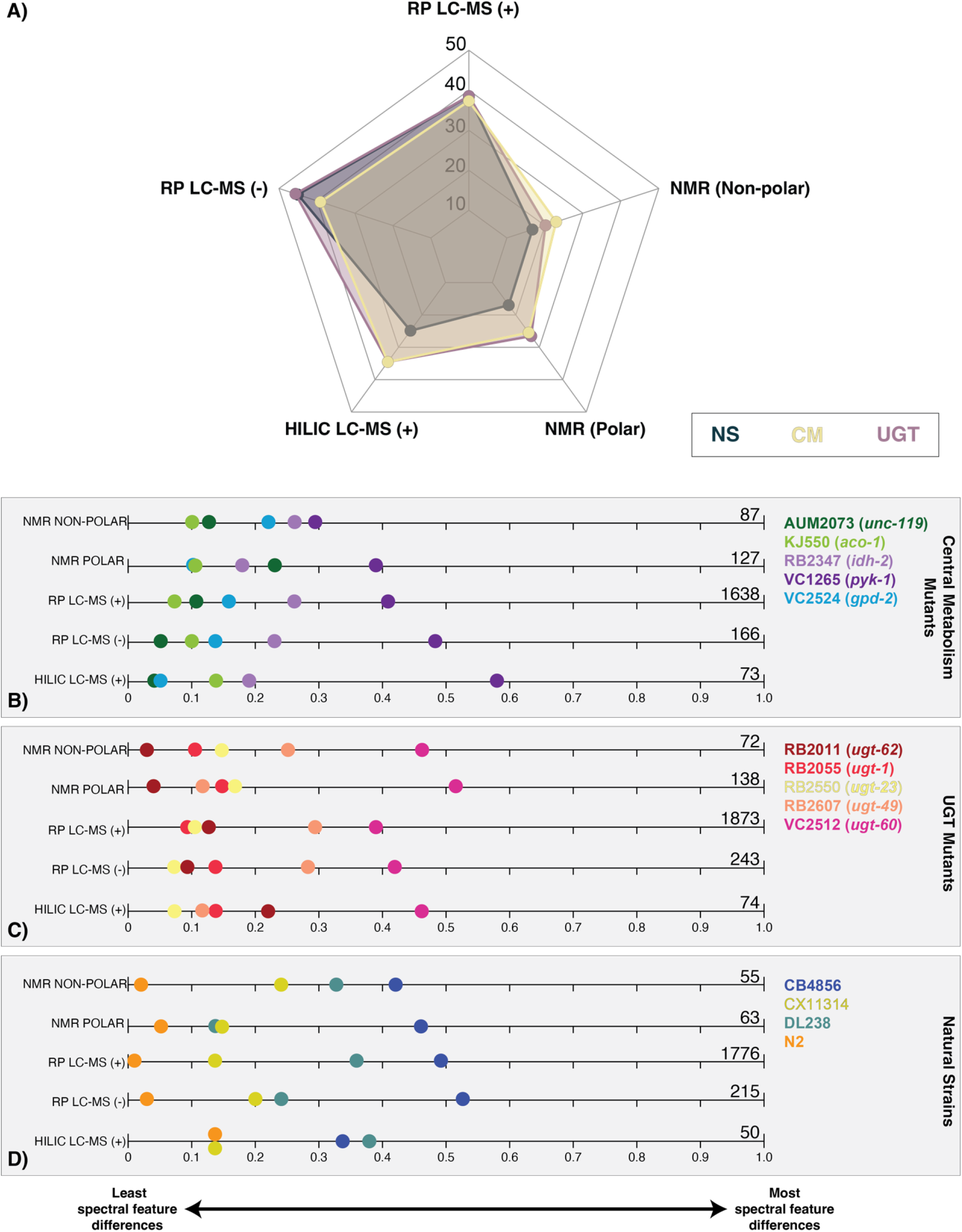
Summary of significant spectral features found in each analytical platform and across the three studies. **(A)** Percent of significant features. The total number of significant features in all strains, by study, is used as the denominator for each of the five technologies. Significant spectral features identified in at least one strain by study are displayed for **(B)** central metabolism mutants, **(C)** UGT mutants, and **(D)** natural strains. Zero indicates the strain has no significant spectral feature differences from PD1074, while one indicates that all spectral feature differences from PD1074 are present in that strain. Significant feature totals are summarized at the end of the plot and detailed in **Table 1**.

**Figure 3.**
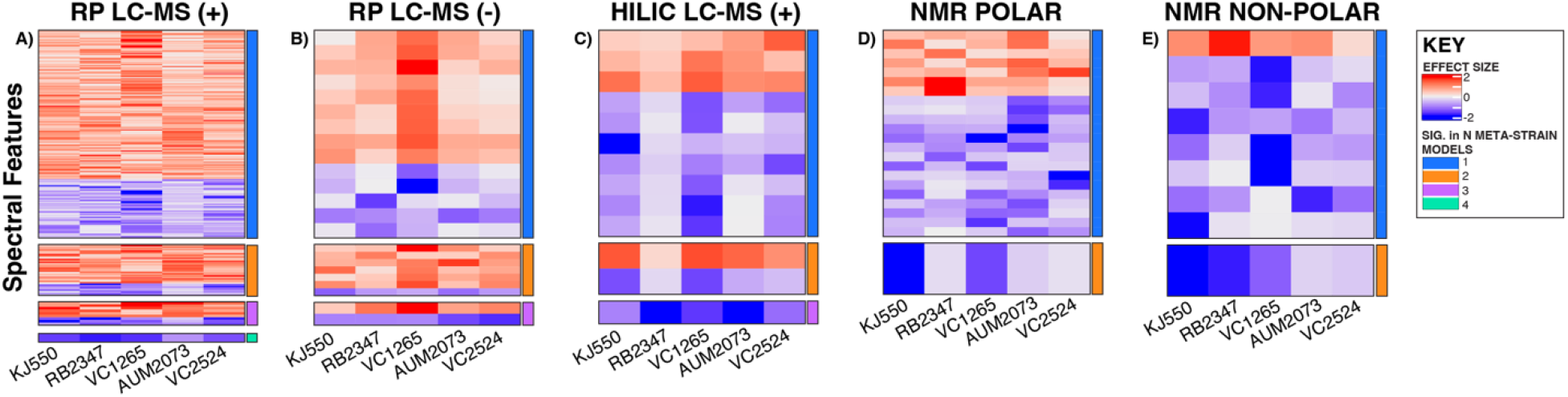
Heatmaps of significant spectral features identified in the CM mutant study. **(A)** RP LC-MS positive mode (**B)** RP LC-MS negative mode (**C)** HILIC LC-MS positive mode (**D)** NMR polar (**E)** NMR non-polar. For each heatmap, the first five columns are strains, and each row represents a spectral feature with an effect size that is consistently higher or lower relative to PD1074 in that study. The effect sizes range from (2 to -2). Positive effect sizes (*i*.*e*., the strain had a higher peak at that given metabolic feature than PD1074) are displayed in red. Negative effect sizes (*i*.*e*., PD1074 had a higher peak at that given metabolic feature than the test strain) are displayed in blue. The right-hand column indicates the number of models in which a given spectral feature is statistically significant. See **Figure S4** for additional CM mutant results.

### Spectral features significant in a mutant and NS

The percentage of significant features in each of the mutant studies (CM and UGT) that overlapped in at least one NS (**Figure 4**) are features of interest for follow-up compound identification. CM mutant strains AUM2073 (*unc-119*) and RB2347 (*idh-2*) share 75% and 68% of their significant features with a NS, respectively. UGT mutants, RB2607 (*ugt-49*) and RB2055 (*ugt-1*) share 67% and 62% of their significant features with a NS, respectively. RB2011 (*ugt-62*) had the most overlap with the NS sharing 67% of its significant features in RP LC-MS (+) and 44% in HILIC LC-MS (+). See **Figures S7 and S8** for significant feature overlap across study comparisons.

**Table 1.** Summary of significant spectral features found in all three studies across NMR and LC-MS. The total number of significant spectral features (*p* < 0.05) for a given strain and each analytical platform are listed.

| Study Group | Strain | Number of significant spectral features |  |  |  |  |
| --- | --- | --- | --- | --- | --- | --- |
|  |  | NMR |  | LC-MS |  |  |
|  |  | Non-polar | Polar | RP + | RP - | HILIC + |
| Central metabolism mutants (CM) | AUM2073 | 11 | 29 | 175 | 8 | 3 |
|  | KJ550 | 9 | 14 | 110 | 17 | 10 |
|  | RB2347 | 23 | 22 | 421 | 38 | 14 |
|  | VC1265 | 25 | 49 | 671 | 79 | 42 |
|  | VC2524 | 19 | 13 | 261 | 24 | 4 |
|  | Total CM sig. features by platform | 87 | 127 | 1638 | 166 | 73 |
| UGT mutants (UGT) | RB2011 | 2 | 6 | 237 | 21 | 16 |
|  | RB2055 | 8 | 20 | 161 | 34 | 10 |
|  | RB2550 | 11 | 23 | 200 | 18 | 5 |
|  | RB2607 | 18 | 17 | 539 | 69 | 9 |
|  | VC2512 | 33 | 72 | 736 | 101 | 34 |
|  | Total UGT sig. features by platform | 72 | 138 | 1873 | 243 | 74 |
| Natural Strains (NS) | N2 | 1 | 3 | 22 | 6 | 7 |
|  | DL238 | 18 | 15 | 631 | 52 | 19 |
|  | CX11314 | 13 | 16 | 254 | 44 | 7 |
|  | CB4856 | 23 | 29 | 869 | 113 | 17 |
|  | Total NS sig. features by platform | 55 | 63 | 1776 | 215 | 50 |
| Total sig. features by platform |  | 146 | 228 | 2541 | 281 | 115 |

**Figure 4.**
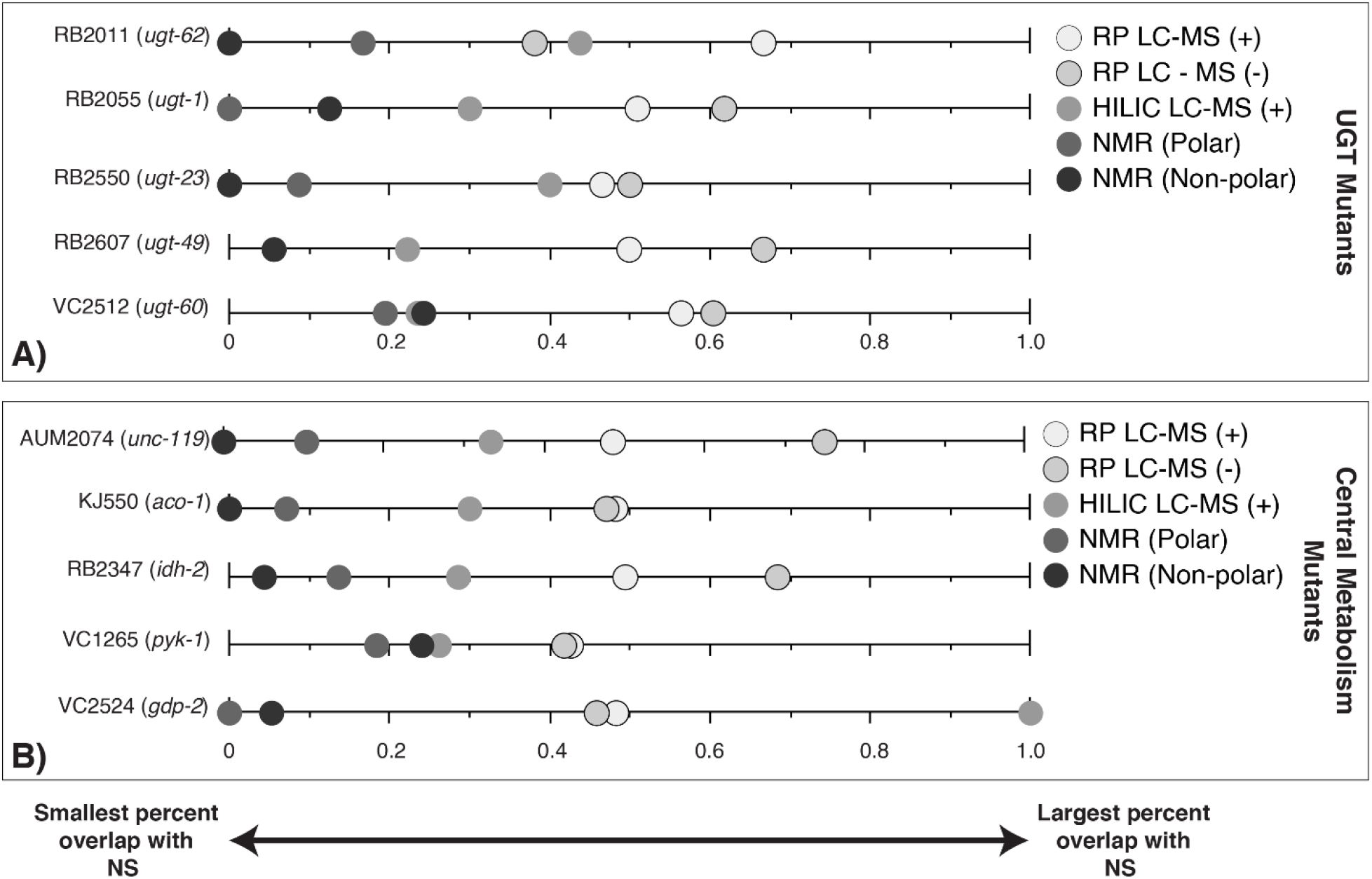
Percent of significant features for each of the mutant studies (CM and UGT) that are also significant in at least one NS by analytical platform. **(A)** UGT mutants **(B)** CM mutants. Data points at zero indicate the analytical platform detected no significant spectral features shared between the mutant strain and a natural strain. Data points at one indicate all significant spectral features for the mutant strain are shared with a natural strain for that analytical platform.

We focused on compounds affected in any CM mutants and used those to identify which UGTs and NS had genetic variation in those same compounds for the NMR polar data. Using COLMAR^61^, we identified three putative compounds significant in strains from all three studies. Of the 35 putative compounds showing evidence for metabolic variation in the NMR data, 13 were annotated (**see Table S2**).

Nine putative compounds show metabolic variation in response to the *pyk-1* mutation (**Figure 5**). The mutation in *pyk-1* affects a large portion of the metabolome. The gene *pyk-1*, is involved in one of the last enzymes of glycolysis, encoding for pyruvate kinase and responsible for glycolytic ATP production. The depletion of lactic acid production is consistent with the mutation in *pyk-1*^62^ in the strain VC1265. We saw the depletion of lactic acid in DL238 (NS), and an increase in VC2512 (*ugt-60*) (**Figure 5**). As expected, none of the 13 compounds identified in the NMR polar dataset were significant in N2 (**Figure 5**). Interestingly, annotated compounds were also similar to PD1074 in CX11314 (NS), RB2055 (*ugt-1*), RB2607 (*ugt-49*), and RB2011 (*ugt-62*).

**Figure 5.**
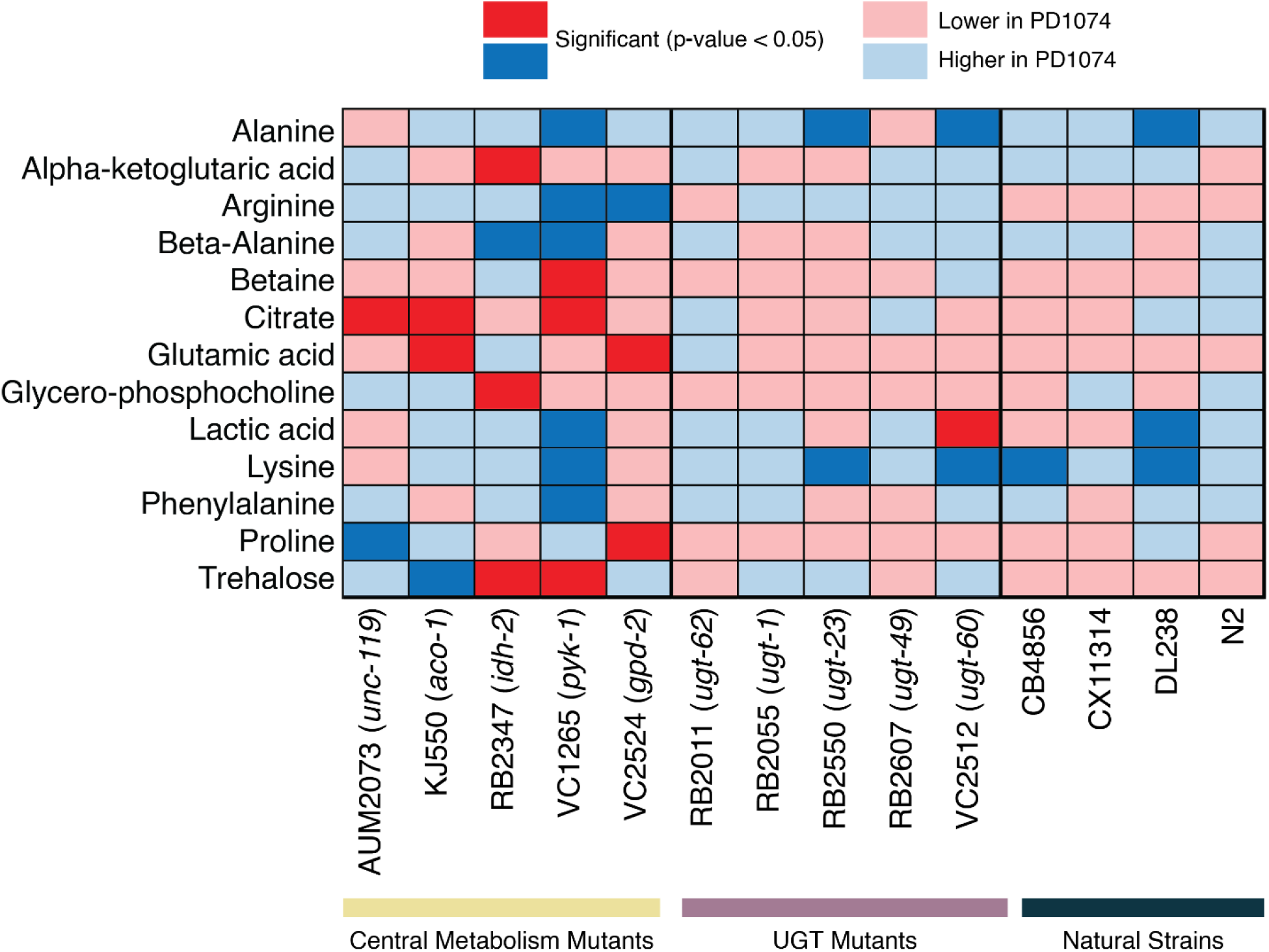
Heatmap of metabolites identified by NMR. Significant NMR spectral features in the central metabolism mutants are compared across UGT mutants and natural strains. Deep blue boxes indicate the metabolite is significant and more abundant in PD1074 compared to the test strain. Deep red boxes indicate the metabolite is significant and more abundant in the test strain. Light colored boxes indicate the direction of effect when the metabolite is not significantly different between PD1074 and the test strain. For compounds with more than one significant feature, the highest effect size feature is used for this figure. The significant compound list provides metabolites to pursue in subsequent experiments. See **Table S2** for compound annotation list and details.

## DISCUSSION

The experimental design enables the same spectral features to be identified across experiments. We elected to use PD1074 to augment the design and focus on stable features in PD1074 instead of attempting to capture unique features in test strains in this analysis. This choice explicitly enables us to use PD1074 to anchor comparisons across studies of genetically diverse test strains, in the current data and in the future. However, the principle behind this approach is not limited to a single strain/genotype but we acknowledge that the inclusion of an augmentation will increase the number of samples, extractions and replicate measurements increasing the cost and size of the study. Individual investigators will need to find their own balance of generalizability and throughput.

A focus on spectral features present despite environmental variation eliminates the need to annotate compounds prior to across study comparisons. The presented technologies can detect many more spectral features that can be identified and then subsequently interpreted biologically. By focusing on stable features, there are far fewer features analyzed here than is typical in untargeted metabolomics. In addition, the analyses presented here fail to identify presence/absence variation, undercount the number of features present, and potentially fail to identify crucial novel metabolites formed as a result of a mutation. However, this is an analytical choice. There is no limitation placed on the technology during data acquisition and these data can also be analyzed for presence/absence variation and for spectral features that vary.

In non-model organism experiments, implementation of a BRM^31, 34^ also enables stable feature identification. New analytic advances have enabled joint alignment and feature selection across high levels of variability when there is a common QC standard like a BRM included in each batch^77^. However, not all experiments have an appropriate BRM. We demonstrate that in the absence of a BRM, pooled anchor (here PD1074) samples can be used to connect the experiment over time and identify stable features even with variable data acquisition conditions.

Perhaps of most value to experimental success is the inclusion of several alternate strategies for stable feature identification. Even though a high abundance of features due to contaminants from sample preparation are present in the LC-MS dataset, the use of individual PD1074 samples along with their pools enable the identification of stable spectral features. This approach is very similar to the BRM approach in the NMR data where both controls are present. Additionally, multiple biological replicates of PD1074 provides the added benefit of enabling a straightforward calculation of the relative effect size for each test strain by batch that can be used later to combine batches or compare test strains statistically using meta-analysis and avoiding a complicated normalization process due to batch variance.

Our analysis focuses on a single anchor strain, PD1074, and allows us to compare the effect of all common stable spectral features across experiments. The same result can be achieved in cohort studies by identifying a control group and including multiple control samples in each batch. Effect sizes for each group relative to the control can be calculated for each batch. Further, if all groups are present in each batch, effect sizes between groups can also be calculated. Spectral features consistently present within groups can be prioritized for future studies so that database matching and, ultimately, compound identification efforts are focused on the most likely biologically important spectral features. This aspect is important as studies^7, 9, 63^ increase in size and complexity^31, 36, 64, 65^.

We demonstrate how a list of significant spectral features can be used to focus NMR compound identification efforts (13 annotated compounds). A similar approach can be used for LC-MS where features are annotated using accurate mass, elemental formula, MS/MS database matching, and *in silico* predictions of spectral features. Compound identification approaches for LC-MS are challenging and oftentimes require orthogonal data for confident annotations. This approach allows future MS/MS experiments to target spectral features that are known to be affected in multiple experiments and are consistently detectable enabling the investigator to collect these specific features rather than relying on the data-dependent acquisition (DDA) or iterative DDA approaches. *In silico* prediction methods for NMR and MS/MS have improved accuracy although ambiguity is expected to remain for large molecular weight formulas^67. 1^H and ^13^C 1D NMR and MS/MS fragmentation *in silico* predictions can be prioritized for target features identified with this approach^66^.

Mapping metabolites in pathways is complicated because many metabolites are involved in multiple pathways and/or have yet to be described. The genetic mutation approach used to annotate gene function in pathways has had limited success in untargeted metabolomics because of the scope of the experiments and the necessity of subsequent rescue experiments to discern pathway-gene relationships. With the large numbers of unknown spectral features, this problem is complex. Meta-analysis allows for the identification of significant spectral features in a straightforward manner when batch effects complicate mixed effects models. Similarly, untargeted studies of collections of genotypes^68^ using an reference genotype, in this case PD1074, can leverage data across experiments and increase the utility of untargeted metabolomics for genetic studies and increase the efficiency of the compound identification process^14^.

## ACKNOWLEDGEMENTS

Research reported in this manuscript was supported by the National Institutes of Health Award Number U2CES030167 (A.S.E., E.C.A., F.E.L., F.M.F., I.J.A., K.M.M., L.M.M.). F.M.F. also acknowledged support from NIG grant 1R01CA218664-01. The authors would like to thank Pamela Kirby at the Edison Lab for assistance with material generation and storage logistics. Thank you to John Glushka for NMR support. Thank you to Oleksandr (Alex) Moskalenko for application and bioinformatics support, and development of SECIMTools 2.0. HiPerGator3, the University of Florida High Performance Computing Platform and it’s support staff, in particular, Erik Deumens for co-ordination. Kelsey Sinclair and Carter Johnson for help with documentation for the SECIMTools 2.0 updates. Some strains were provided by the *Caenorhabditis* Genetics Center (CGC), which is funded by NIH Office of Research Infrastructure Programs (P40 OD010440). Lastly, we thank WormBase.

## AUTHOR CONTRIBUTIONS

Conceptualization, L.M.M. and A.S.E.; Methodology, A.O.S., B.M.G., A.M.M., and L.M.M.; Software, L.M.M. and A.M.M.; Validation, A.O.S., B.M.G., Z.L., R.M.B., A.M.M., and L.M.M.; Formal Analysis, A.O.S., B.M.G, A.M.M., and L.M.M.; Investigation, A.O.S., B.M.G., G.J.G., and C.A.K.; Resources, A.O.S., B.M.G, G.J.G., and C.A.K.; Data Curation, A.O.S., B.M.G., G.J.G., A.M.M., and L.M.M.; Writing – Original Draft, A.O.S. and B.M.G.; Writing – Review &Editing, A.O.S., B.M.G., G.J.G., C.A.K., R.M.B., A.M.M. E.C.A., F.M.F., A.S.E., and L.M.M.; Visualization, A.O.S., B.M.G, G.J.G., A.M.M., and L.M.M; Supervision, A.M.M., F.E.L. III, E.C.A., I.J.A., F.M.F, A.S.E., and L.M.M.; Funding Acquisition, A.S.E.

## METHODS

### C. elegans Strain Selection

This study used 15 *Caenorhabditis elegans* strains obtained from the *Caenorhabditis* Genetics Center (CGC) and *Caenorhabditis elegans* Natural Diversity Resource (CeNDR) ^41^. Fourteen *C. elegans* strains were used as ‘comparison strains,’ and one strain, PD1074, was used as the ‘reference strain’ (**Table S1**). These strains were selected to cover the diversity of interests in the metabolomics community, to encompass samples with mutations in primary and secondary metabolism, along with natural strains.

### C. elegans Sample Growth and Preparation

Large populations of nematodes were generated for every biological replicate with minimal variability^43^. The stable *Escherichia coli* IBAT BRM and food source used throughout this experiment was described previously^31^. Briefly, a large-scale culture plate (LSCP) was used for each biological sample to generate a large mixed-stage population of worms (four to seven LSCP replicates per test strain). For each LSCP, worms were collected, population size estimated, and subsequently divided into at least 12 identical aliquots of 200,000 worms in ddH_2_O and flash-frozen in liquid nitrogen to quench metabolism and stored at -80°C^43^. As a QC sample, *C. elegans* IBAT BRM was generated and saved in 200,000 worm aliquots^31^.

### Study Design

Each *C. elegans* strain was reared with and harvested with at least one PD1074 LCSP. Sample collection for all three studies lasted more than six months. To ensure handling was consistent, no more than five LSCPs were handled at a given time. There are 29 independent PD1074 LCSPs collected and 104 independent test strain sample LSCPs. The PD1074 represents an augmented design^37, 39^, where one PD1074 biological replicate (‘check’) was matched with each test strain biological replicate (‘new treatments’).

### Iterative Batch Average Method (IBAT) in PD1074

An IBAT control^31^, made up of pools of PD1074, was generated to assess batch variance across the six batches in this study. Briefly, aliquots of PD1074 were pooled together to generate a BRM that (i) minimizes the variance between batches of PD1074 BRM, (ii) can be used throughout large-scale experiments, and (iii) can be used to determine the magnitude of variation at multiple points in a metabolomics experiment. See Gouveia *et al*., 2021 for more details on the IBAT process^31^.

### Lyophilization

Frozen aliquots of 200,000 *C. elegans* worms were retrieved from -80°C and lyophilized in a VirTis® BenchTop™ “K” Series Freeze Dryer (*SP Industries, Inc*.). After lyophilization, each aliquot was weighed and stored at -80°C until homogenization.

### Batching and Quality Control Across Analytical Platforms

Up to 24 extractions could be performed simultaneously based on centrifuge capacity limitations. Six extraction batches were needed to accommodate all the strains. Extraction batches were designed in sets of two consecutive batches so that each test strain has all replicates measured in close proximity. The test strains belong to three studies, and the studies are used to create three sets of two batches each, for a total of six batches. The three sets were collected back-to-back in NMR but are separated in time by some months in the LC-MS, although the column and instrument are the same for all three sets. There was a needle failure between batches 5 and 6 in the HILIC LC-MS run. The NS were collected in batches 1 and 2, most of the CM mutants in batches 3 and 4 (exception, AUM2073 and VC2524 were collected in batches 5 and 6), and the UGT mutants were mostly in batches 5 and 6 with (exception, RB2011 was collected in batch 1). Each extraction batch includes half of the replicates for each test sample type (balanced across two consecutive batches), a set of PD1074 LCSPs, the IBAT control, and an extraction blank. Extraction blanks were processed with test strain and PD1074 aliquots to control for homogenization and extraction steps to account for non-biologically related LC-MS or NMR features that arise from sample preparation. Test LSCPs were unique to a batch, but aliquots from the same PD1074 LSCPs may be included more than once. Multiple aliquots of the same LSCP enables QC of feature selection and alignment were included as these differ only by technical variance (*i*.*e*., instrument and extraction) (**Figure 1**).

### NMR Sample Homogenization and Extraction

Frozen lyophilized *C. elegans* aliquots were retrieved from -80°C. 200 μL of 1 mm zirconia beads (BioSpec Products) were added to each sample and homogenized at 420 rcf for 90 seconds in a FastPrep-96 homogenizer and subsequently placed on dry ice for 90 seconds to avoid overheating; this step was repeated twice for a total of three rounds.

Using the homogenized samples, 1 mL of 100% IPA chilled to -20°C was added to the lyophilized/homogenized sample powder and Zirconia beads in two increments of 500 μL. After each addition of 500 μL, samples were vortexed for 30 seconds – 1 min., and left at room temperature (RT) for 15 - 20 minutes. After RT incubation, samples were stored overnight (∼12 hours) at -20°C. Samples were centrifuged for 30 minutes at 4°C (20,800 rcf). The supernatant was transferred to a new tube to analyze non-polar molecules. 1 mL of pre-chilled 80:20 CH_3_OH:H_2_O (4°C) was added to the remaining worm pellet to analyze polar molecules. The polar fraction was allowed to shake at 4°C for 30 minutes. Samples were centrifuged at 20,800 rcf for 30 minutes at 4°C. The supernatant was transferred to a new tube to analyze non-polar molecules. Both polar and non-polar samples were placed in a Labconco Centrivap at RT and monitored until completely dry. Once dry, polar samples were reconstituted in D_2_O (99%, Cambridge Isotope Laboratories, Inc.) in a 100 mM sodium phosphate buffered solution with 0.11 mM sodium 2,2-dimethyl-2-silapentane-5-sulfonate (DSS-D6; 98%; Cambridge Isotope Laboratories, Inc.). Once dry, non-polar samples were reconstituted in CDCl_3_ (99.96%; Cambridge Isotope Laboratories, Inc.). Samples were vortexed until fully soluble, and 45 μL of each sample were transferred into 1.7 mm NMR tubes (Bruker SampleJet) for acquisition.

### NMR Acquisition

To collect the polar fraction, one-dimensional (1D) ^1^H NMR spectra were acquired with a noesypr1d pulse sequence on a NEO 800 MHz Bruker NMR spectrometer equipped with a 1.7mm TCI cryoprobe and a Bruker SampleJet autosampler cooled to 6°C. During acquisition, 32,768 complex data points were collected using 128 scans with two additional dummy scans. The spectral width was set to 15 ppm.

To collect the non-polar fraction, one-dimensional (1D) ^1^H NMR spectra were acquired with a zg pulse sequence (zg30). During acquisition, 65,536 complex data points were collected using 64 scans with four additional dummy scans. The spectral width was set to 20.2 ppm.

In addition, immediately after each 1D acquisition, a 2D J-resolved spectrum is collected using the Bruker pulse program jresgpprqf. For both the polar and non-polar fractions, 8,192 and 40 points were collected using eight scans, four dummy scans, and spectral widths of 16 and 0.09 ppm, respectively. See metabolomics workbench Study IDs (NMR polar: ST002095; NMR non-polar: ST002096) for additional acquisition parameters and data.

For metabolite identification the web server COLMARm was used. As inputs three two-dimensional experiments 1H-1H TOCSY (dipsi2gppphzspr), 1H-13C HSQC (hsqcetgpsisp2.2) and 1H-13C HSQC-TOCSY (hsqcdietgpsisp.2) collected on separate pooled PD1074 polar samples were used. The HSQC experiment was collected using 6250 and 720 points in the indirect and direct dimensions, 32 scans and 16 dummy scans and a spectral width of 13 ppm for the proton and 165 ppm for the carbon dimensions. The HSQC-TOCSY experiment parameters were identical to HSQC except for 32 dummy scans and a 90 ms mixing time. The TOCSY experiment was collected with 7272 points and 800 points in the indirect and direct dimensions, 32 scans and 16 dummy scans, a spectral width of 11.367 ppm in both dimensions and a mixing time of 90 ms. Peak picking and spectral match against hydrophilic metabolite databases (*i*.*e*., HMDB and BMRB) was carried out by COLMARm using 0.04 and 0.3 ppm chemical shift cutoffs for ^1^H and ^13^C respectively and a matching ratio cutoff of 0.6. See metabolomics workbench Study IDs (NMR polar: ST002095; NMR non-polar: ST002096) for all the acquisition parameters and data.

### NMR Data Processing

Following data acquisition, the data were processed using NMRPipe ^69^. Fourier transform, an exponential line broadening of 1.5 Hz and manual phase correction were carried out. Using the tools from (MATLAB, The MathWorks, R2019a^70^), the spectra were referenced at 7.24 ppm using the CDCl_3_ resonance, and the polar extracts are referenced at 0.00 ppm using DSS. Solvent regions were removed followed by baseline correction using a statistical smoothing function^71^. Alignment was performed using CCOW^72^ and PAFFT^73^ algorithms. Manual curation of semi-automated peak-picking was carried out by peak picking that used a binning algorithm^74^ to extract peak heights. This was done separately for blanks and samples. Individual spectral features were removed if detected in the solvent and process blanks using the BFF function in SECIMtools^75^.

Two-dimensional NMR experiments were also processed using NMRPipe. Spectra were Fourier transformed, a 90° shifted sine window function and automatic zero filled applied, manually phased and referenced to DSS.

Stable spectral features were compared between individual PD1074 samples, PD1074 pools, and IBAT controls.

### LC-MS Sample Homogenization and Extraction

Using glass and zirconium oxide beads, the aliquots were homogenized for three minutes in a Qiagen Tissuelyser 2. Homogenized worms were extracted with 1.5 mL of isopropanol (IPA) at -20°C overnight (approximately 12 hours), then pelleted and the supernatant transferred to separate 2 mL centrifuge tubes. Supernatants were then dried to completion in a Labconco Centrivap and stored at -80°C for non-polar LC-MS analysis. The pellet was extracted a second time using 80:20 methanol:water (CH_3_OH:H_2_O) (v:v) for 20 minutes at RT while shaking at 1500 rpm. Samples were again pelleted to separate proteins, and the supernatant was transferred to separate 2 mL centrifuge tubes, dried down to completion, and stored at -80°C for polar LC-MS analysis.

### LC-MS Acquisition and Processing

Each instrument run for a single batch included the following controls with replicate injections at the beginning and end of the batch: instrument control, extraction blanks, pooled test sample aliquots, and pooled PD1074 sample aliquots. In the middle of the batch, individual test samples and PD1074 samples were injected in a randomized order.

Non-polar extracts were reconstituted in 75 µL of IPA containing isotopically labeled lipid standards and analyzed by LC-MS using a ThermoFisher Scientific Accucore C30 150 × 2.1mm, 2.6 µm column paired with a Thermo Fisher Orbitrap ID-X in positive and negative polarity. Polar (80:20 CH_3_OH:H_2_O) extracts were reconstituted in 75 µL of 80:20 CH_3_OH:H_2_O containing isotopically labeled arginine, hypoxanthine, hippuric acid, and methionine (Cambridge Isotope Laboratories, Inc.) and analyzed by LC-MS using a Waters BEH Amide 150 × 2.1 mm, 1.7 µm column paired with a Thermo Fisher Orbitrap ID-X in positive and negative polarity. LC-MS/MS data for each mode of analysis was collected using three rounds of iterative DDA (Thermo Scientific AcquireX) performed on pooled test samples.

Data for each sample was collected in full MS1 with a resolution of 240,000 FWHM (full-width half-maximum) and MS/MS spectra of pooled samples were collected at a resolution of 30,000 FWHM using a 0.8da isolation window and stepped HCD collision energies of 15, 30, and 45. See supplemental information for detailed LC-MS parameter settings. Thermo .raw files were converted to centroid mode and .mzML format using Proteowizard’s MSconvertGUI tool^76^. Raw files are deposited at metabolomics workbench Study ID ST002092. Pre-processing steps, input parameters, and set values used for LC-MS data are listed in **Table S4**.

### Selection of stable LC-MS spectral features

A plasticizer contamination event precluded us from quantitatively assessing the performance of an IBAT control in the LC-MS experiments. Instead, we used the 12 PD1074 pools in a two-step procedure. First, the PD1074 pools were averaged over extraction variance for the batch, capturing instrumentation variation across batches. In the second step, we retained the subset of peaks only present in 100% of the individual PD1074 samples to focus on stable peaks across growth conditions (environmental variation). Here, we focus on peaks present across multiple individual samples of the same genotype, PD1074, a variant of the laboratory-adapted strain N2, but any strain of *C. elegans* could serve this purpose.

The 12 pooled PD1074 samples were used to estimate optimal parameters, and then these parameters were applied to all samples using a memory-efficient algorithm SLAW (https://github.com/zamboni-lab/SLAW).^77^ Only spectral features above the blank threshold of 100 for all 12 PD1074 pools were retained for further analysis. SLAW offers the following peak picking algorithms: XCMS centWave^78, 79^, OpenMS FeatureFinderMetabo^80, 81^, and MZmine ADAP^82, 83^ For this study, ADAP was selected.^84^. The SLAW algorithm is predicated upon the assumption that the experimental design includes identical QC samples across an experiment (*e*.*g*., BRM) in intervals during data collection. This inclusion is typical in large-scale studies^31, 85, 86^, but the selection of stable spectral features across extraction variance is not standard. While the benefits of including QC samples are known and recently have been implemented in peak picking and alignment optimization workflows that traditionally have not scaled to large data^87^, the inclusion of PD1074 replicate samples during sample generation, analytical measurement, and data processing is novel.

Spectral peaks were filtered further using the individual PD1074 samples. We took a conservative approach requiring 100% of the PD1074 samples to have each spectral feature present above the blank. This focuses the experiment and our attention on spectral features that are likely to be present in a subsequent independently prepared MS2 experiment in the compound identification process, and not spectral features present sporadically due to variation in growth or extraction.

### Quality Control Assessments for LC-MS and NMR Data

Stable spectral features were rank transformed (*i*.*e*., raw data is replaced by ranks where the lowest rank has the smallest peak height, and the highest rank has the largest peak height for a given spectral feature). QC assessments included Standard Euclidean Distance (SED), principal component analysis (PCA), coefficient of variation (CV), Bland Altman (BA), and sample density distributions to identify potential feature artifacts and/or atypical samples^75^. See **Table S3** for QC parameters and thresholds used to identify stable mass features^75^. PCA is used to visualize distortions due to batch or genotype. BA plots on pools and PD1074 samples within a batch were used to visualize alignment variation, and BA plots on replicate aliquots of the same PD1074 samples were used to verify the success of the alignment across batches. Per feature CV is examined to identify any wildly aberrant features and was used to help refine the quantification of the solvent front.

Sample outliers were identified based on the SED plots. Chromatograms of samples whose distance to other samples did not cross the 95% percentile for the distribution of pairwise distances were manually examined for chromatography failure. The PD1074 LSCP sample “aos54” failed the QC assessment for NMR. The PD1074 LSCP samples “aos53” and “aos41” failed the QC assessment for RP LC-MS datasets. Test strain “aos49” in batch 5 is removed from all datasets, and test strain “aos25” in batch 1 was removed from the HILIC LC-MS positive dataset. Samples were removed from further consideration in their respective datasets.

### Meta-analysis on LC-MS and NMR Data

All replicates of a particular genotype were contained within two sequential batches: however, different test strains within the same study span multiple sets. We used meta-analysis for each feature to compare the test genotype to the control, where each batch is treated as an ‘experiment’ using a fixed effects (FE) model using standardized mean difference (SMD)^75^, referred to as “meta-strain” model throughout. Positive effect sizes indicate that the test strain has a higher peak than PD1074 for a given chemical feature. Negative effect sizes indicate PD1074 has a higher peak than the test strain for a given chemical feature. Each strain was tested against PD1074 to see if that feature is differentially expressed between PD1074 and the test strain. We also used meta-analysis to compare test genotypes to each other, referred to as the “meta-study” model throughout. For example, for the five UGT mutants (*i*.*e*., RB2011, RB2550, RB2055, RB2607, and VC2512) we tested whether a feature is differentially expressed between the test strain and PD1074 in all five test genotypes.

### NMR ^1^H 1D spectra Annotation

Significant features obtained from the meta analysis of the CM mutants were selected for identification. The 2D experiments HSQC, HSQC-TOCSY, and TOCSY were collected from a pooled PD1074 sample. These data served as inputs to the public webserver COLMARm^88^ (Complex Mixture Analysis by NMR), an application that allows us to simultaneously and interactively compare multiple 2D spectra data to HMDB^89^, BMRB^90^, and NMRShiftDB^91^ publicly available databases. Only the significant features were annotated. Annotation confidence scores per compound are detailed in **Table S2** according to the previously reported levels as described elsewhere^92^.

Further annotation details can be found in the COLMAR outputs submitted to Metabolomics Workbench. **Figure 5** illustrates the annotated compounds. Only the feature with the highest effect size was selected for compounds with more than one significant feature. After a list of compounds was identified, WormFlux^44^ was used to explore the effects of the CM mutants on the *C. elegans* metabolic network.

## Code Availability

The python code for QA/QC is available through GitHub (https://github.com/secimTools/SECIMTools) and can be run via a Galaxy install (https://docs.galaxyproject.org/en/master/) or from a command line interface. The meta-analysis (meta_analysis.py) and rank transformation (add_group_rank.py) python code are available on the SECIMtools GitHub page. The Matlab functions used as well as instructions and version control are available at: https://github.com/artedison/Edison_Lab_Shared_Metabolomics_UGA.

## Data Availability

These data are available at the NIH Common Fund’s National Metabolomics Data Repository (NMDR) website, the Metabolomics Workbench, **https://www.metabolomicsworkbench.org** where it has been assigned Study ID (LC-MS: ST002092; NMR polar: ST002095; NMR non-polar: ST002096). The data can be accessed directly via its: Project DOI: http://dx.doi.org/10.21228/M82978. This work is supported by NIH grant U2C-DK119886. Methods and protocols used in this study are available on protocols.io: NMR: dx.doi.org/10.17504/protocols.io.b2rbqd2n LCMS: (dx.doi.org/10.17504/protocols.io.bahjib4n).

