## Supplemental Information for "An anchored experimental design and meta-analysis approach to address batch effects in large-scale metabolomics"

##### *Reverse Phase (RP) chromatography method*

Non-polar extracts were separated using a Vanquish liquid chromatograph (ThermoFisher Scientific), fitted with a ThermoFisher Scientific Accucore™ C30 UPLC RP column (2.1 x 150 mm, 2.6 µm particle size). The compounds were eluted with the following gradient: 60:40 acetonitrile:water (ACN:$H_2O$) with 10 mM ammonium formate and 0.1% formic acid (mobile phase A) and 90:10 isopropanol:acetonitrile with 10 mM ammonium formate and 0.1% formic acid (mobile phase B) using the following gradient program: 0.0 min 20% B; 1.0 min 60% B; 5.0 min 70% B; 5.5 min 85% B; 8.0 min 90% B; 8.2-10.5 min 100% B; 10.7-12.0 min 20% B. A curve 5 value was set for 0.0 minutes, and a curve 6 for the remainder of the gradient. The flow rate was set at 0.400 mL $min^{-1}$. The column temperature was set to 50°C, and the injection volume was 2 µL. The following internal standards were spiked in for RP analysis: 15:0-18:1(d7) PC, 15:0-18:1(d7) PE, 15:0-, 18:1(d7) PS, 15:0-18:1(d7) PG, 15:0-18:1(d7) PI, 18:1(d7) LPC, 18:1(d7) LPE, 18:1(d7) Chol Ester, 15:0-18:1(d7) DG, 15:0-18:1(d7)-15:0 TG, 18:1(d9) SM, and Cholesterol (d7).

##### *Hydrophilic Interaction Liquid Chromatography (HILIC) method*

Polar extracts were separated using a Vanquish liquid chromatograph (ThermoFisher Scientific), fitted with a Waters Acquity UPLC BEH Amide column (2.1 x 150 mm, 1.7 µm particle size). The compounds were eluted with the following gradient: 80:20 water:acetonitrile ($H_2O$:ACN) with 10 mM ammonium formate and 0.1% formic acid (mobile phase A) and 100% ACN with 0.1% formic acid (mobile phase B) using the following gradient program: 0.0-0.5 min 95% B; 8.0-9.4 min 40% B; 9.5-11.0 min 95% B. A curve 5 value was set for 0.0 minutes, a curve 6 at 0.5 min, curve 7 at 8.0 min, and a curve 6 for the remainder of the gradient. The flow rate was set at 0.400 mL $min^{-1}$. The column temperature was set to 40°C, and the injection volume was 2 µL.

##### Mass spectrometer settings and methods

An Orbitrap ID-X Tribrid mass spectrometer (ThermoFisher Scientific) equipped with a HESI ion source was used for all mass spectrometry data collection. For HILIC analysis the mass spectrometer was run in full MS mode at a resolution of 240,000 (FWHM at *m/z* 200) for the duration of the chromatographic gradient. A normalized automatic gain control (AGC) target of 100% was set with a maximum injection time of 100 ms. A tune file with the following source conditions was used for positive and negative mode: spray voltage (+) 3500, spray volage (-) 2500, vaporizer temperature: 275 ºC, sheath gas: 40, aux gas: 8, sweep gas: 1, and S-Lens RF level: 60%. Scan range covered *m/z* 70-1050. Calibration was conducted using ThermoFisher Pierce™ Negative Ion Calibration Solution and Pierce™ LTQ Velos ESI Positive Ion Calibration Solution prior to collecting negative and positive mode data, respectively.

Identical parameters were used for reverse-phase analysis with the following exceptions: spray voltage (+) 3500, spray volage (-) 2800, maximum injection time of 200ms, vaporizer temperature: 425 ºC, sheath gas: 60, aux gas: 18, sweep gas: 4, and a scan range of *m/z* 150-2000.

##### *Growth of C. elegans using the LSCP method yields on average 2.4 million mixed-stage worms per sample*

Growth of *C. elegans* using the LSCP method generates large mixed-stage populations of *C. elegans* with minor handling and manipulation of the animals, ideal for metabolomics experiments. Population dynamics depend on the strain's behavior (*i.e.*, burrowing strains tend to have lower worm recovery) and growth success (*i.e.,* contamination). The LSCP method yields population sizes from approximately 94,500 to 9,290,000. The mean population size within the refernce strain, PD1074, and across strains is approximately 2.4 million worms (**Figure S2**). PD1074 LSCPs take between 10 – 14 days to grow to a full mixed-stage population. The mean growth time for PD1074 is ten days. The slowest growing strain grows for a maximum of 20 days, and the fastest growing strain for a minimum of 10 days (**Figure S2**). Strain-dependent variation in growth rate is to be expected because of the wide range of strains and traits included in this study. Notably, growth time is not indicative of the final population size (**Figure S2**). Additional information on *C. elegans* growth and population dynamics has been presented previously[43].

##### *Variation between PD1074 and N2*

The reference strain PD1074, obtained from CeNDR, is a traceable variant of the laboratory-adapted N2 Bristol strain[93]. Here, we include N2, obtained from CeNDR, as one of our natural strains as an additional way to validate our methods and processes. Phenotypically, compared to PD1074, N2 is not significantly different in the amount of time needed to cultivate a population or the resulting population size (**Figure S2**). Metabolically, N2 is most similar to PD1074 showing 16 total feature differences in the RP LC-MS (+) data, six feature differences in the RP LC-MS (-), and seven differences in the HILIC LC-MS (+) data (**Table 1**), indicating little difference between N2 and PD1074. In the non-polar NMR data, N2 has three significant feature differences and seven significant feature differences in the polar NMR data (**Table 1**), having many fewer spectral features differentiating from PD1074. Both the phenotypic and metabolic data showcase that these strains are very similar. In contrast, in the RP LC-MS modes, N2 has up to a 50-fold difference from the other strains (**Figure S5**).

***Model comparison for batch effect corrections***

Meta-analysis is a statistical analysis that combines summary statistics instead of an analysis of individual samples[60, 94]. A meta-analysis can be used to account for the batch effects in untargeted metabolomics. In a traditional meta-analysis, an effect size is calculated for each study and then combined and weighed by the individual study sample sizes [95, 96]. As meta-analysis is a promising approach to address the complicated variance structure in a straightforward way, and has been shown to be equivalent to more complex linear model approaches on individual data on larger sample sizes[94], we demonstrate that meta-analysis, even with relatively small sample sizes per group (n=6 for test samples), is very similar to a mixed effects model with the variance modeled appropriately[55]. This study design makes it possible to apply both approaches and compare inferences directly. In other more complex situations modeling the individual-level data can be very challenging. The meta-analysis model was formally compared to the linear model in several formulations, and in each case, the meta-strain model has similar statistical inferences as the linear model, consistent with the literature on larger sample sizes[60]. To mirror the comparison in Lin & Zeng (2010)[94], a fixed-effect model or random effect model can be used to infer a true biological effect from the effects estimated from individual batches. The fixed-effect model assumes that the underlying effect sizes in all batches are identical and treats the inverse of the variance of each batch effect size as the weight to account for all variabilities in each batch.

Contrary to the fixed-effect model, a random-effect model assumes that the underlying effect size in all batches is similar but not identical. As there are just a few batches in this study, a fixed-effect model is applied. We use the fixed-effect model effect size estimate:

$$\bar{\theta}_w = \frac{\sum_{i=1}^{k} w_i \, \theta_i}{\sum_{i=1}^{k} w_i}$$

where $w_i$ is a weight calculated as the inverse of variance for the effect size in batch *i*, and $\bar{\theta}_w$ is the effect size of interest inferred from the individual effect sizes in batches. For the ANOVA comparison the effect size is calculated as:

$$\bar{\theta} = \frac{lsmean_{test} - \; lsmean_{PD1074}}{sd}$$

$$sd = \sqrt{n} * se$$

To illustrate the comparability between these approaches, we compare the linear model by batch *l*, where strain *i* is the independent variable and ion signal for each spectral feature *m,* and test replicate *j* is the response variable*:*

$$Y_{mlij} = \mu + batch_l + \; strain_i + e_{lij}$$

The example depicted is typical, with the number of features detected in the linear model consistently slightly higher than the meta-analysis[60]. This is consistent with the slight benefit in the degrees of freedom estimates from the combined model on individual data. The effect size estimates are consistent. Batch effects across experiments are an enormous problem in metabolomics experiments, and the inability to adequately address this in a mixed model analyses is a well-known problem[55, 60]. Meta-analysis presents a straightforward way of comparing data across experiments.

#### SUPPLEMENTAL FIGURES

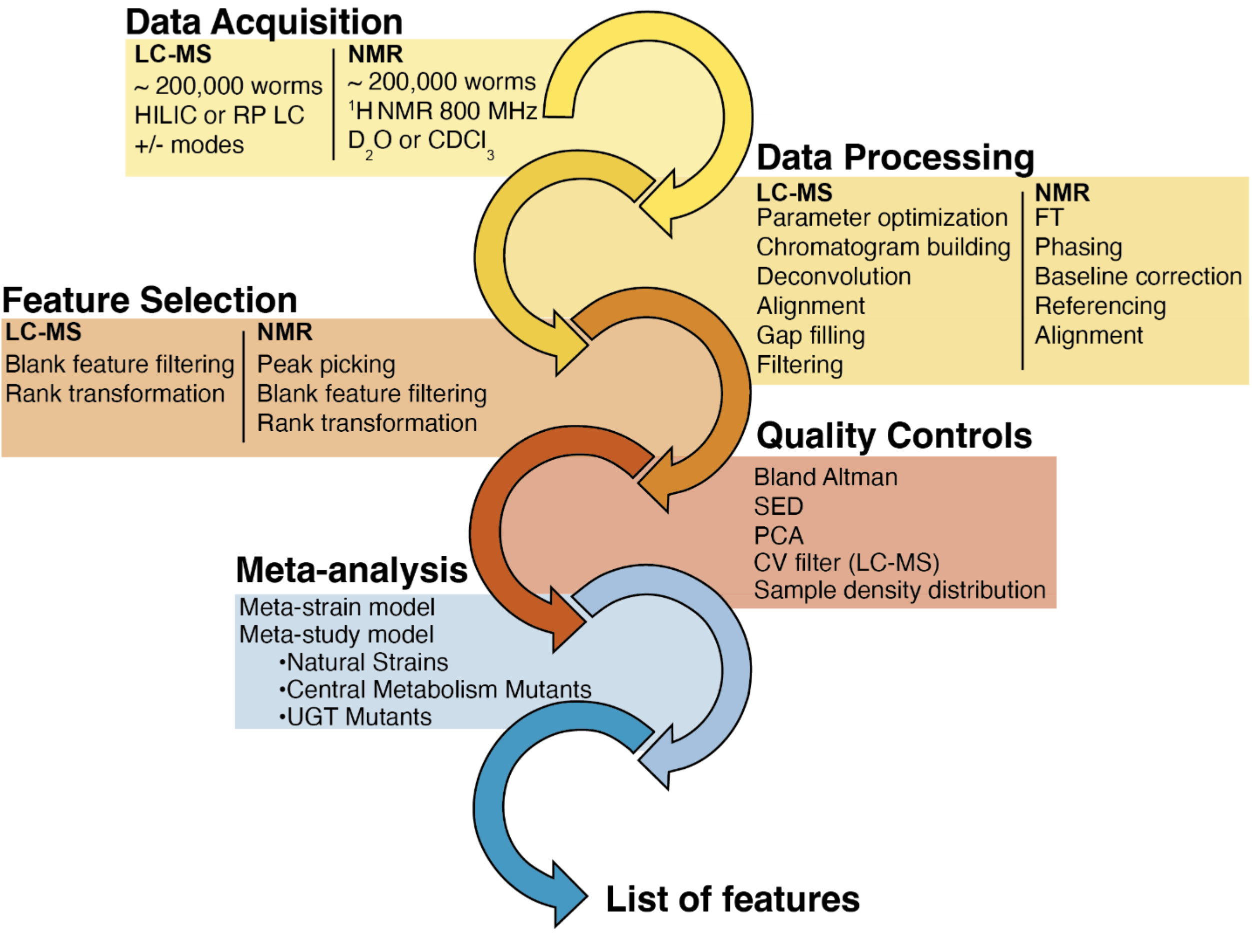

**Supplemental Figure 1. Unified workflow for spectral feature selection across platforms.** Steps and software listed to obtain spectral features. LC-MS-specific steps are listed on the left side of each process. NMR-specific steps are denoted on the right side of each process. When neither NMR nor LC-MS is indicated, the same steps are performed on data from both analytical platforms (see Methods for detailed documentation).

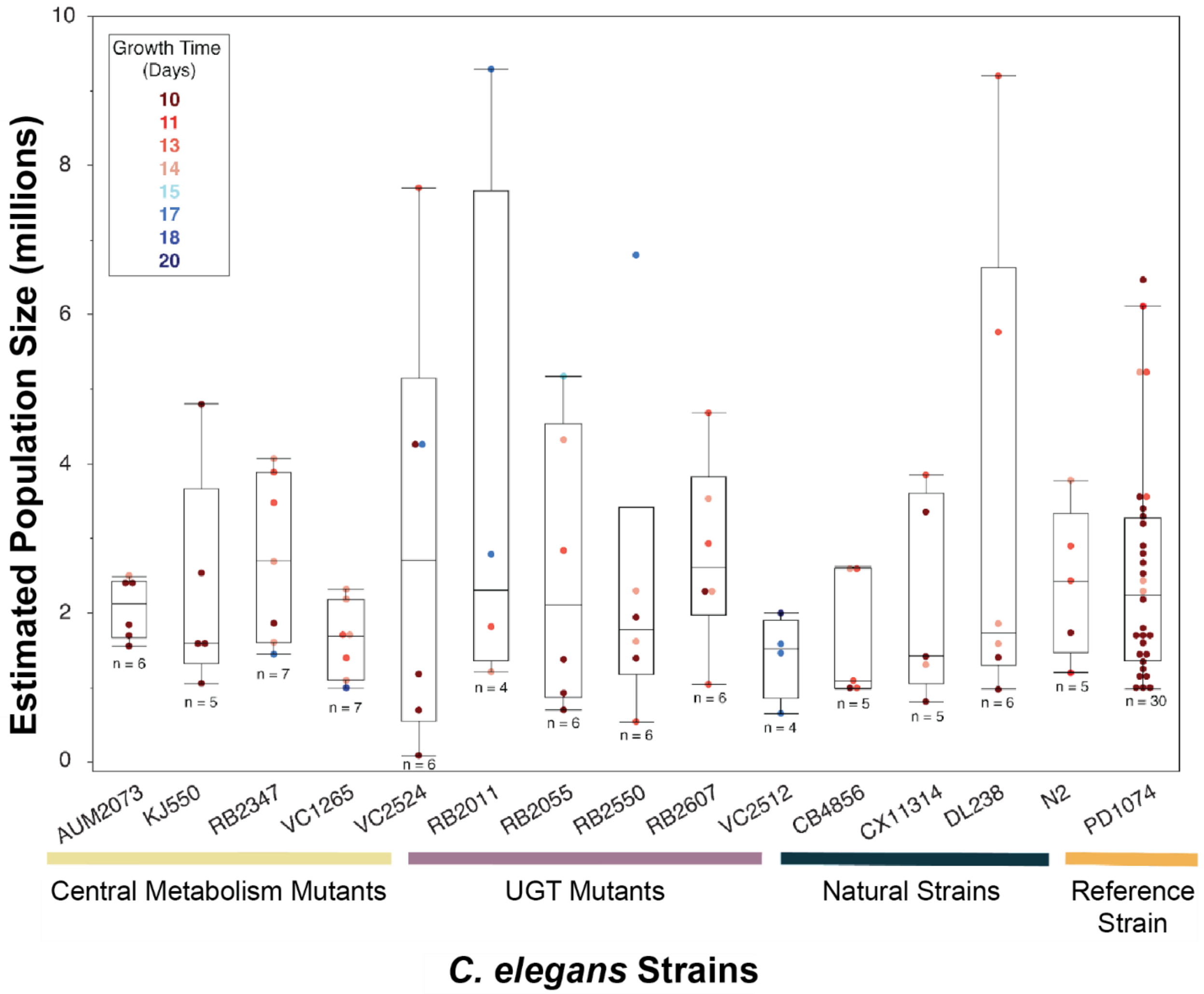

**Supplemental Figure 2**. **LSCP method generates, on average, a population of 2.4 million mixed-stage *C. elegans* populations.** The LSCP yields population sizes in the smallest population growths at around 94,500 worms and the biggest population growths at around 9,290,000 worms. The mean population size across all strains was 2.4 million worms. Bars underneath *C. elegans* strain names indicate each strains study. Comparisons of population size for all pairs using Tukey's HSD test were performed. No significant differences are observed between estimated population sizes across *C. elegans* strains. Colored data points indicate the growth time (days) to generate a given LSCP sample.

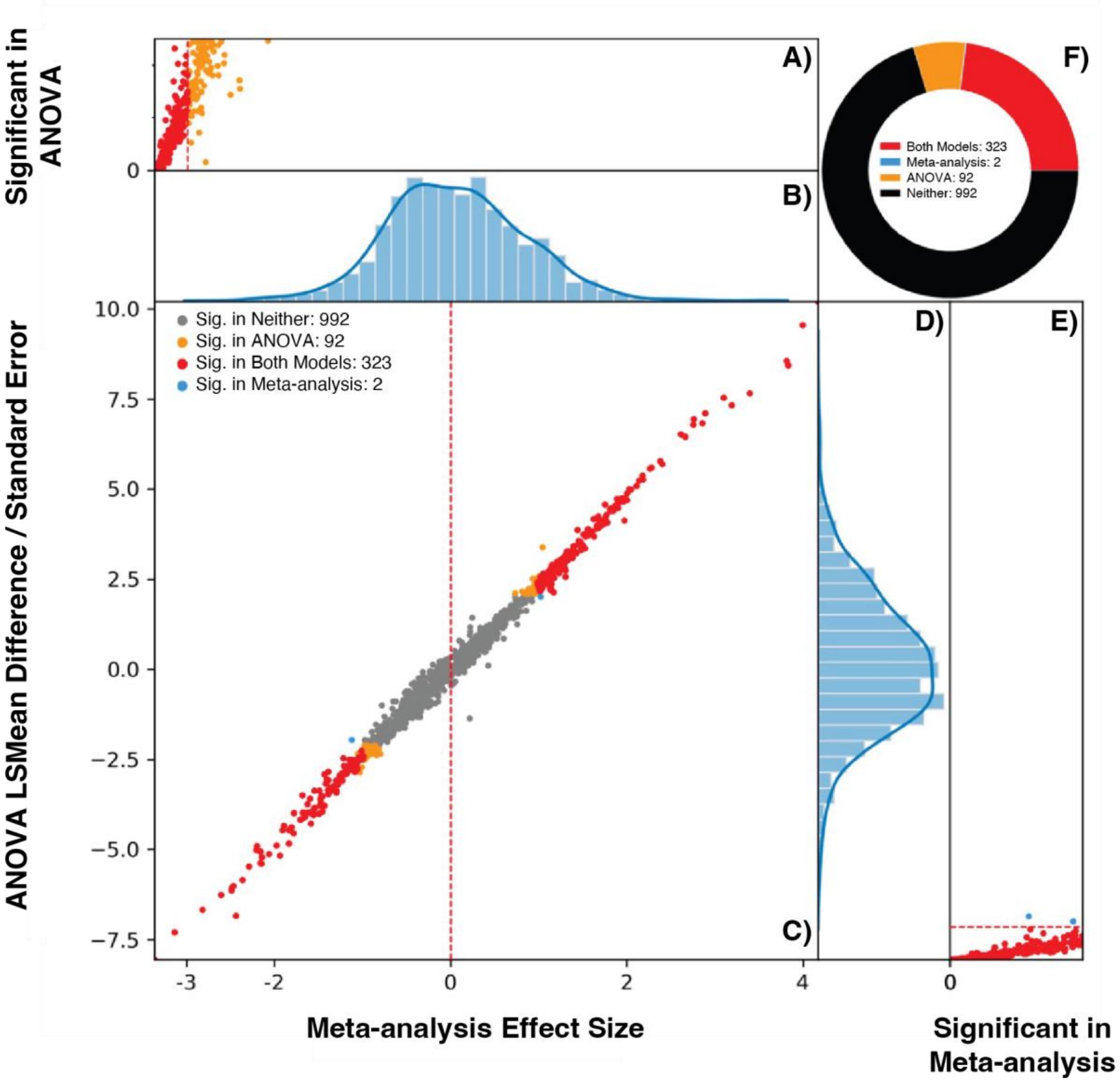

**Supplemental Figure 3: Comparing the inferences from a linear model and meta-analysis using the test genotype, VC1265.** There are six panels in the plot. **(A)** Displays the p-values where the y-axis is from 0-1 and represents the significant results (p-value < 0.05) in the linear model. **(B)** displays the effect size distribution for effect sizes in the x-axis for the scatter plot. **(C)** is a scatter plot, where the x-axis is the effect sizes calculated by the meta-anlaysis model and the y-axis is the *lsmean* difference calculation. Each point is one spectral feature. **(D)** displays the effect size distribution for effect sizes in the y-axis for the scatter plot. Point colors represent significance of the test of the null hypothesis where the mean peak height for VC1265 is not different from the mean peak height of PD1074 with a nominal threshold p-value < 0.05. Red points are significant in both models. Orange points are significant in the linear model. Blue points are significant results in the meta-analysis model. Grey points indicates that spectral feature is not significant in either model. Red lines in **(A)** and **(E)** are significance thresholds for the nominal p-value < 0.05 on the other test. **(F)** summarizes the results.

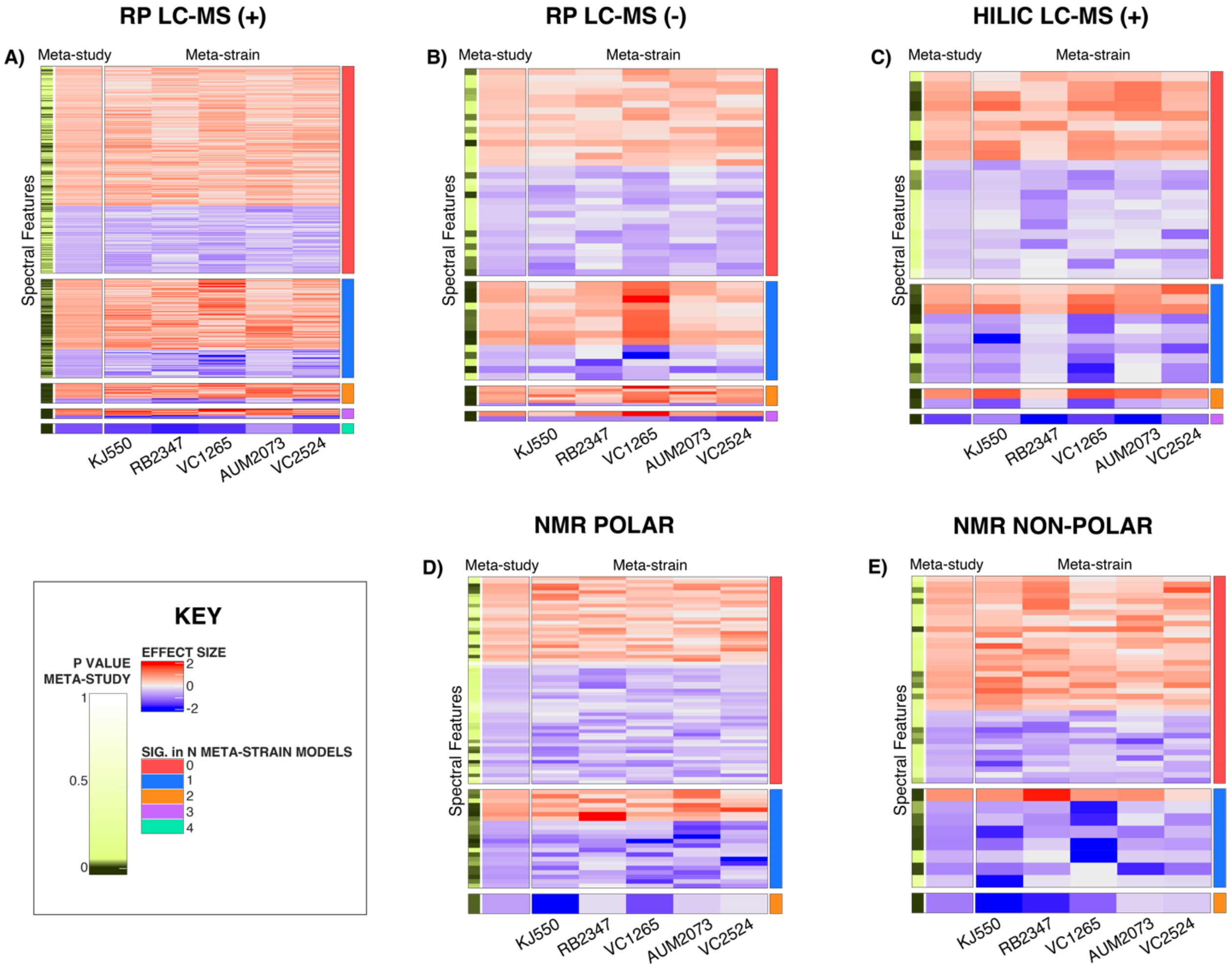

**Supplemental Figure 4. Heatmaps of significant spectral features identified via the meta-strain and meta-study models in the CM mutants study. (A)** RP LC-MS positive mode **(B)** RP LC-MS negative mode **(C)** HILIC LC-MS positive mode **(D)** NMR polar and **(E)** NMR non-polar. The first two columns on the left-hand side pertain to the meta-study model results. The yellow and black bar highlights the significant features found in the meta-study model, followed by the meta-study heatmap. The following five columns within the heatmap compare the spectral features for a given strain in the meta-strain model. For each heatmap, each row represents a spectral feature with an effect size that is consistently higher or lower relative to PD1074 in that study (meta-strain) or across strains (meta-study). The effect sizes range from (2 to -2). Positive effect sizes (*i.e.*, the strain had a higher peak at that given metabolic feature than PD1074) are displayed in red. Negative effect sizes (*i.e.,* PD1074 had a higher peak at that given metabolic feature than the test strain) are displayed in blue. The right-hand column indicates the number of strains in which a given spectral feature is statistically significant in the meta-strain model.

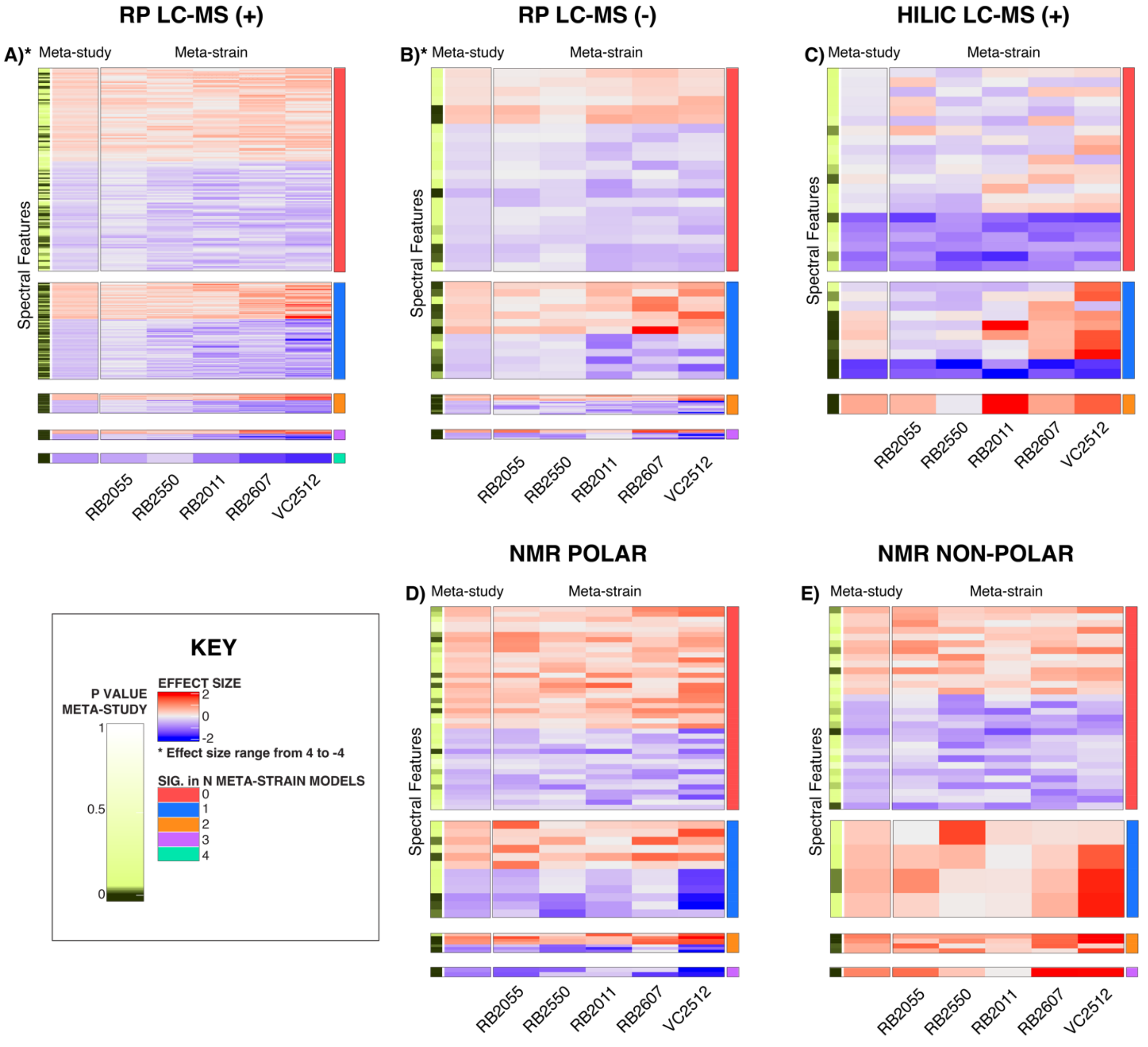

**Supplemental Figure 5. Heatmaps of significant spectral features identified via the meta-strain and meta-study models in the UGT study. (A)** RP LC-MS positive mode **(B)** RP LC-MS negative mode **(C)** HILIC LC-MS positive mode **(D)** NMR polar and **(E)** NMR non-polar. The first two columns on the left-hand side pertain to the meta-study model results. The yellow and black bar highlights the significant features found in the meta-study model, followed by the meta-study heatmap. The following five columns within the heatmap compare the spectral features for a given strain in the meta-strain model. For each heatmap, each row represents a spectral feature with an effect size that is consistently higher or lower relative to PD1074 in that study (meta-strain) or across strains (meta-study). The effect sizes range from (2 to -2). Modes denoted with an asterisk have effect sizes that range from (4 to -4). Positive effect sizes (*i.e.*, the strain had a higher peak at that given metabolic feature than PD1074) are displayed in red. Negative effect sizes (*i.e.*, PD1074 had a higher peak at that given metabolic feature than the test strain) are displayed in blue. The right-hand column indicates the number of strains in which a given spectral feature is statistically significant in the meta-strain model.

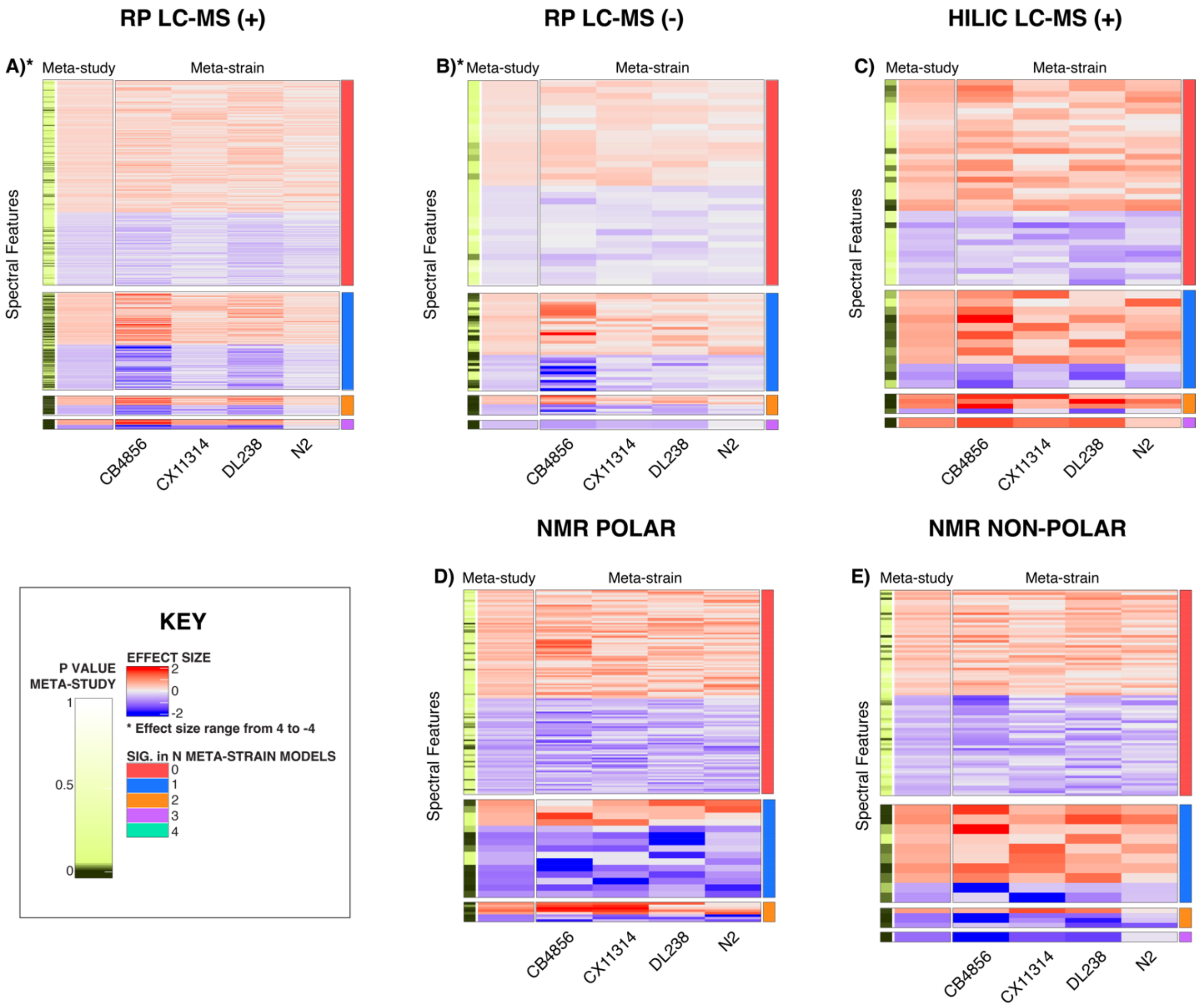

**Supplemental Figure 6. Heatmaps of significant spectral features identified via the meta-strain and meta-study models in the NS study. (A)** RP LC-MS positive mode **(B)** RP LC-MS negative mode **(C)** HILIC LC-MS positive mode **(D)** NMR polar and **(E)** NMR non-polar. The first two columns on the left-hand side pertain to the meta-study model results. The yellow and black bar highlights the significant features found in the meta-study model, followed by the meta-study heatmap. The following five columns within the heatmap compare the spectral features for a given strain in the meta-strain model. For each heatmap, each row represents a spectral feature with an effect size that is consistently higher or lower relative to PD1074 in that study (meta-strain) or across strains (meta-study). The effect sizes range from (2 to -2). Modes denoted with an asterisk have effect sizes that range from (4 to -4). Positive effect sizes (*i.e.*, the strain had a higher peak at that given metabolic feature than t PD1074) are displayed in red. Negative effect sizes (*i.e.*, PD1074 had a higher peak at that given metabolic feature than the test strain) are displayed in blue. The right-hand column indicates the number of strains in which a given spectral feature is statistically significant in the meta- strain model.

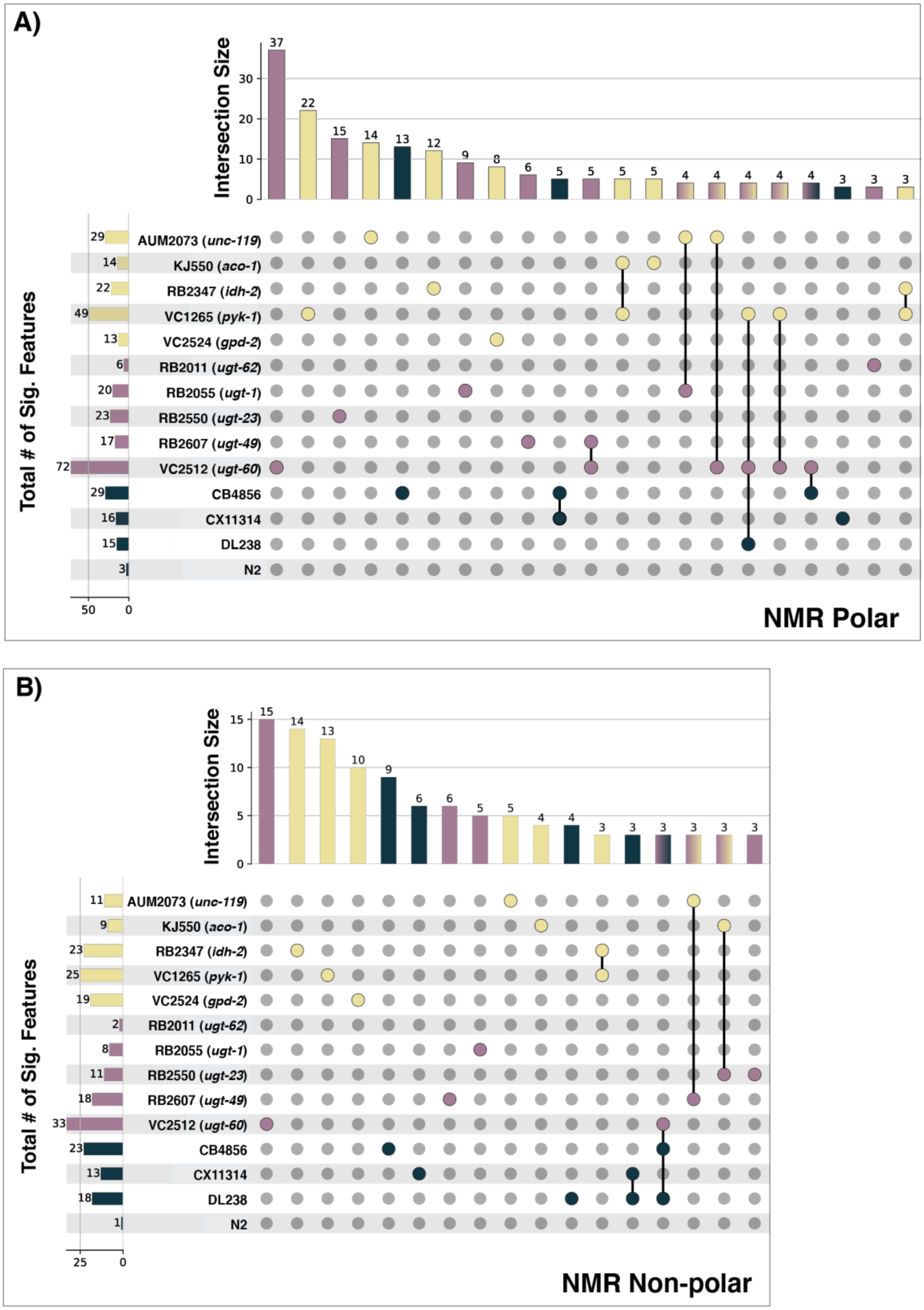

**Supplemental Figure 7. Significant features found via the meta-strain model across the three study groups in NMR.** (**A)** NMR polar (**B)** NMR non-polar. Fourteen strains in the CM mutants (yellow), UGT mutants (purple), and NS (green) are displayed. Horizontal bar plots sum the total number of significant features for a strain. Vertical bar plots sum significant feature interactions within and across strains. Significant feature connections below three are not displayed.

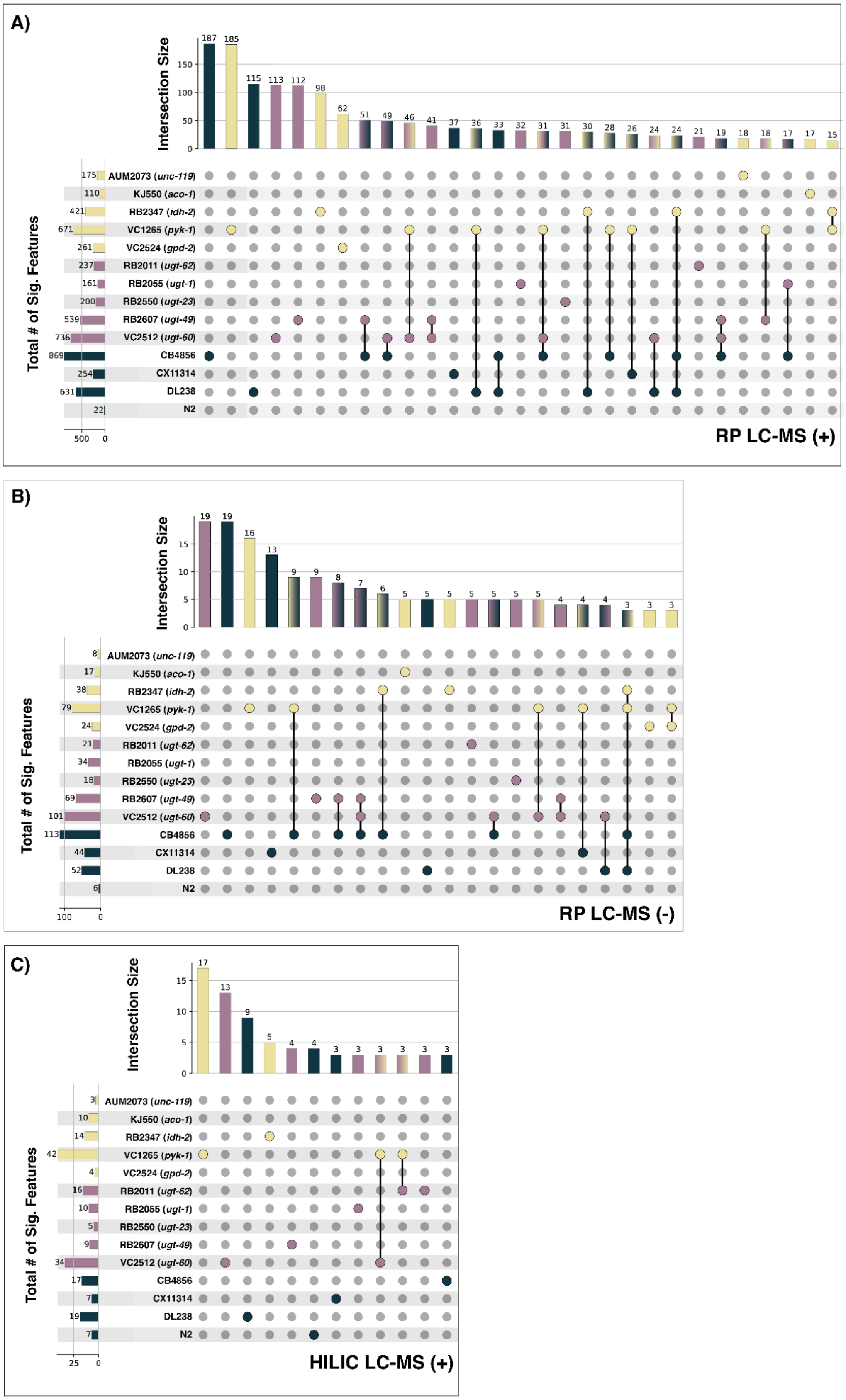
A)
Intersection Size
Total # of Sig. Features
AUM2073 (unc-119)
KJ550 (aco-1)
RB2347 (idh-2)
VC1265 (pyk-1)
VC2524 (gpd-2)
RB2011 (ugt-62)
RB2055 (ugt-1)
RB2550 (ugt-23)
RB2607 (ugt-49)
VC2512 (ugt-60)
CB4856
CX11314
DL238
N2
RP LC-MS (+)
B)
RP LC-MS (-)
C)
HILIC LC-MS (+)

**Supplemental Figure 8. Significant features found via the meta-strain model across the three study groups in LC-MS. (A)** RP LC-MS positive mode (**B)** RP LC-MS negative mode (**C)** HILIC LC-MS positive mode. Fourteen strains in the CM mutants (yellow), UGT mutants (purple), and NS (green) are displayed. Horizontal bar plots sum the total number of significant features for a strain. Vertical bar plots sum significant feature interactions within and across strains. Significant feature connections below three are not displayed for all modes, except for the RP LC-MS positive mode, where connections below 15 are not displayed.

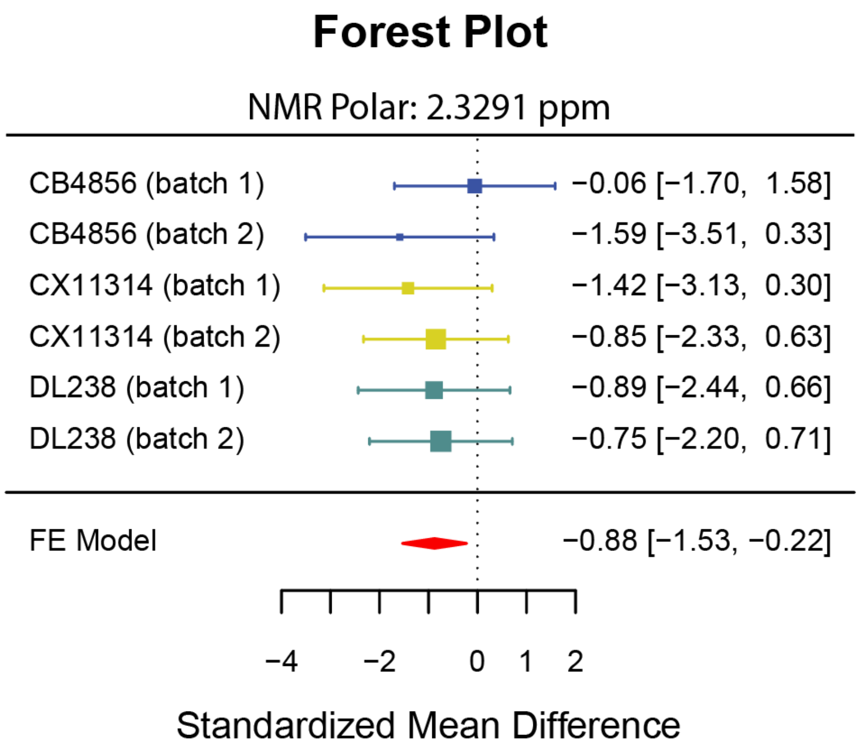

**Supplemental Figure 9. Forest plot comparing the meta-strain and meta-study models in the NS for the NMR polar ppm 2.3291.** The meta-strain model for each NS in each batch is displayed, where the area of each square is proportional to the study's weight in the meta-analysis with confidence intervals (CI) represented by whiskers. The dashed vertical line represents the overall measure of effect. The right-hand column is the measure of effect (odds ratio) for each study. The meta-study model (FE model) is represented by the red diamond, where the lateral points indicate CI for the estimate.

### SUPPLEMENTAL TABLES

| Strain | Genotype | Source | About | Study Group |
|---|---|---|---|---|
| PD1074 | wild type | CeNDR | wild type | Anchor Strain |
| AUM2073 | *unc-119; vizSi34 II; unc-119(ed3) III.* | CGC | GSK-3 promotes S-phase entry and progress in germline cells to maintain tissue output | Central Metabolism Mutants |
| KJ550 | *aco-1 (jh131) X.* | | Exhibits aconitate hydratase activity. Is involved in tricarboxylic acid metabolic process. Localizes to cytosol. Is expressed in seam cell. Is an ortholog of human ACO1 (aconitase 1). | |
| RB2128 | *idh-1; F59B8.2(ok2832) IV.* | | *idh-1 encodes a predicted cytosclic isocitrate dehydrogenase; by homology, IDH-1 is the formation of alpha-ketoglutarate from isocitrate, Is predicted to have isocitrate dehydrogenase (NADP+) activity and metal ion binding activity. Is expressed in tail. Is an ortholog of human IDH1 (isocitrate dehydrogenase (NADP(+)) 1).* | |
| RB2347 | *idh-2(ok3183) X.* | | Is predicted to have isocitrate dehydrogenase (NADP+) activity and metal ion binding activity. Human ortholog(s) of this gene are implicated in D-2-hydroxyglutaric aciduria 2. Is an ortholog of human IDH2 (isocitrate dehydrogenase (NADP(+)) 2). | |
| VC1265 | *pyk-1; F25H5.3(ok1754) I.* | | *pyk-1 encodes for one of two pyruvate kinases. essential for embryonic development, Is predicted to have kinase activity; metal ion binding activity; and pyruvate kinase activity. Human ortholog(s) of this gene are implicated in pyruvate kinase deficiency of red cells. Is an ortholog of human PKLR (pyruvate kinase L/R) and PKM (pyruvate kinase M1/2).* | |
| VC2524 | *gpd-2 (ok3243) X.* | | Is predicted to have NAD binding activity; NADP binding activity; and glyceraldehyde-3-phosphate dehydrogenase (NAD+) (phosphorylating) activity. Is expressed in several structures, including AB; Psub1; and head. Is an ortholog of human GAPDH (glyceraldehyde-3-phosphate dehydrogenase). | |
| RB2011 | *ugt-62 (ok2663)* | CGC | homozygous. gene knockout ugt-62, Is predicted to have glucuronosyltransferase activity. Human ortholog(s) of this gene are implicated in Crigler-Najjar syndrome and Gilbert syndrome. Is an ortholog of several human genes including UGT1A6 (UDP glucuronosyltransferase family 1 member A6); UGT1A8 (UDP glucuronosyltransferase family 1 member A8); and UGT1A9 (UDP glucuronosyltransferase family 1 member A9). | UGT Mutants |
| RB2055 | *ugt-1 (ok2718) V.* | | homozygous. gene knockout ugt-1, Is predicted to have UDP-glycosyltransferase activity. Human ortholog(s) of this gene are implicated in Crigler-Najjar syndrome and Gilbert syndrome. Is an ortholog of several human genes including UGT1A4 (UDP glucuronosyltransferase family 1 member A4); UGT1A8 (UDP glucuronosyltransferase family 1 member A8); and UGT1A9 (UDP glucuronosyltransferase family 1 member A9). | |
| RB2550 | *ugt-23 (ok3541) X.* | | homozygous. gene knockout ugt-23, Is predicted to have glucuronosyltransferase activity. Is involved in gastrulation. Human ortholog(s) of this gene are implicated in Crigler-Najjar syndrome and Gilbert syndrome. Is an ortholog of human UGT3A1 (UDP glycosyltransferase family 3 member A1) and UGT3A2 (UDP glycosyltransferase family 3 member A2). | |
| RB2607 | *ugt-49(ok3633) V.* | | homozygous. gene knockout ugt-49, Is predicted to have glucuronosyltransferase activity. Human ortholog(s) of this gene are implicated in Crigler-Najjar syndrome and Gilbert syndrome. Is an ortholog of several human genes including UGT2A3 (UDP glucuronosyltransferase family 2 member A3); UGT2B10 (UDP glucuronosyltransferase family 2 member B10); and UGT2B11 (UDP glucuronosyltransferase family 2 member B11). | |
| VC2512 | *ugt-60(ok3248) III/hT2 [bli-4(e937) let-?(q782) qIs48] (I;III).* | | homozygous. gene knockout ugt-60, Is predicted to have glucuronosyltransferase activity. Human ortholog(s) of this gene are implicated in Crigler-Najjar syndrome and Gilbert syndrome. Is an ortholog of human UGT2B7 (UDP glucuronosyltransferase family 2 member B7). | |
| N2 | wild type | CeNDR | wild type | Natural Strains |
| DL238 | | | | |
| CX11314 | | | | |
| CB4856 | | | | |

**Supplemental Table 1.** ***C. elegans*** **genotypes used in study.**

| Feature ID | Putative Annotation | Strain | Confidence score |
|---|---|---|---|
| ppm_1_4761 | Alanine | VC1265 | 4 |
| ppm_1_4896 | | | |
| ppm_2_5027 | Alpha-ketoglutaricacid | RB2347 | 3 |
| ppm_2_237 | Aminoadipic Acid* | RB2347 | 2 |
| ppm_3_2462 | Arginine | VC1265 | 4 |
| ppm_3_2547 | | VC2524 | |
| ppm_1_6361 | Arginine* | AUM2073 | 4 |
| ppm_1_6525 | | | |
| ppm_3_1839 | Beta-Alanine | RB2347 | 4 |
| ppm_3_2672 | Betaine | VC1265 | 4 |
| ppm_0_86209 | CH3-Lipoprotein | AUM2073 | 1 |
| ppm_2_5584 | Citrate (carbon shift outside threshold criteria) | AUM2073 | 2 |
| ppm_2_5788 | | KJ550 | |
| ppm_2_5866 | | AUM2073 | |
| ppm_2_0689 | Glutamic Acid | VC2524 | 4 |
| ppm_2_0778 | | | |
| ppm_2_1094 | | KJ550 | |
| ppm_2_1186 | | | |
| ppm_2_3474 | Glutamic Acid* | RB2347 | 4 |
| ppm_2_3585 | | | |
| ppm_2_4498 | Glutamine-Unknown* | AUM2073 | 4 |
| ppm_2_4587 | | | |
| ppm_2_4673 | | | |
| ppm_2_4774 | | | |
| ppm_2_1285 | Glutamine/Glutamic Acid* | RB2347 | 3 |
| ppm_3_5782 | Glycerol* | AUM2073 | 3 |
| ppm_3_9436 | Glycero-phosphocholine | RB2347 | 3 |
| ppm_1_3238 | Lactic Acid | VC1265 | 4 |
| ppm_1_3396 | | | |
| ppm_1_6686 | Leucine* | AUM2073 | 4 |
| ppm_1_755 | Lysine | KJ550 | 4 |
| ppm_3_0189 | | VC1265 | |
| ppm_3_029 | | | |
| ppm_3_0393 | | | |
| ppm_1_9208 | Lysine-Acetic Acid-Arginine* | AUM2073 | 3 |
| ppm_1_6832 | Lysine-Arginine* | KJ550 | 4 |
| ppm_1_7156 | | VC1265 | |
| ppm_1_7348 | | | |
| ppm_7_3231 | Phenylalanine | VC1265 | 4 |
| ppm_7_338 | | | |
| ppm_7_4387 | | RB2347 | |
| ppm_3_1231 | Phenylalanine* | RB2347 | 4 |
| ppm_7_3876 | | | |
| ppm_2_007 | Proline (low level) | AUM2073 | 3 |
| ppm_2_0306 | | VC2524 | |
| ppm_3_4544 | Trehalose | RB2347 | 4 |
| ppm_3_4704 | | VC1265 | |
| ppm_5_188 | | RB2347 | |
| ppm_5_2069 | | KJ550 | |
| ppm_3_6408 | Trehalose-Glycerol* | VC1265 | 4 |
| ppm_3_6575 | | RB2347 | |
| ppm_3_8518 | Trehalose* | KJ550 | 4 |
| ppm_3_8699 | | | |
| ppm_1_591 | Unknown 1 | AUM2073 | n/a |
| ppm_1_5981 | | | |
| ppm_1_6058 | Unknown 2 | KJ550 | n/a |
| ppm_3_2244 | Unknown 3 | VC1265 | n/a |
| ppm_8_1863 | Unknown 4 | AUM2073 | n/a |
| ppm_0_94007 | Unknown 5* | AUM2073 | n/a |
| ppm_1_9545 | Unknown 6* | KJ550 | n/a |
| ppm_2_2768 | Unknown 7* | RB2347 | n/a |
| ppm_2_2952 | Unknown 8* | RB2347 | n/a |
| ppm_3_7919 | Unknown 9* | RB2347 | n/a |

**Supplemental Table 2.** List of significant features and respective annotation. FeatureID indicates the chemical shift of each feature. Putative annotation indicates compound name as obtained from COLMAR. Strain indicates the corresponding mutant for each feature deemed significantly different (p-value < 0.005). Confidence score defined as 1 to 5, with 5 being the highest. The scale is defined as follows: (1) putatively characterized compound classes or annotated compounds, (2) matched to literature and/or 1D spectra of a reference standard, (3) matched to HSQC, (4) matched to HSQC and validated by HSQC–TOCSY and TOCSY

(COLMARm), and (5) validated by spiking the authentic compound into the sample[92]. Unknown has no matches in COLMAR database. Low level indicates features were low intensity in 2D spectra. Abbreviations: asterisk – feature indicated was found to be overlapped within the 2D spectra, n/a – not applicable.

| SECIM Tools Workflow: | Input Parameters | Set Value |
|---|---|---|
| Blank Feature Filtering (BFF) | BFF Threshold | 5000 |
| | Criterion Value | 100 |
| Standard Euclidean Distance (SED) | Group/Treatment [Optional] | genotype |
| | Input Run Order Name [Optional] | run_order |
| | Additional groups to separate by [Optional] | |
| | Threshold | 0.95 |
| Coefficient of Variation (CV) | Group/Treatment [Optional] | |
| | CV Cutoff [Optional] | 0.1 |
| Principal Components Analysis (PCA) | Group/Treatment [Optional] | |
| Bland-Altman (BA) | Outlier Cutoff | 3 |
| | Sample Flag Cutoff | 0.2 |
| | Feature Flag Cutoff | 0.05 |
| | Group/Treatment [Optional] | genotype |
| | Group Name [Optional] | |
| Generate distribution of features across samples | Group/Treatment [Optional] | genotype |

**Supplemental Table 3.** SECIM Tools workflow, input parameters, and set values used for QC steps on analytical data.

| | RP (+) | HILIC (+) | RP (-) |
|---|---|---|---|
| fold blank | 1 | 1 | 1 |
| frac qc | 1 | 1 | 1 |
| alpha* | 0.1 | 0.1 | 0.1 |
| dmz* | 0.005 | 0.005 | 0.005 |
| drt* | 0.03 | 0.03 | 0.03 |
| extracted quantity | height | height | height |
| num references | 150 | 150 | 150 |
| ppm | 5 | 5 | 5 |
| adducts positive | [M+H]+, [M+2H]2+, [M+Na]+, [M+K]+, [M+NH4]+, [M+2Na-H]+, [2M+H]+, [2M+2H]2+, [2M+H+Na]2+, [2M+Na]+, [2M+2Na-H]+, [M+2H-NH3]2+, [M+H-H2O]+, [M+H-H2O]+, [M+2H-H2O]2+, [M+3H]3+, [M+CH3COONa+H]+, [M+CH3COONa+Na]+, [M+CH3COONa+NH4]+ | [M+H]+, [M+2H]2+, [M+Na]+, [M+K]+, [M+NH4]+, [M+2Na-H]+, [2M+H]+, [2M+2H]2+, [2M+H+Na]2+, [2M+Na]+, [2M+2Na-H]+, [M+2H-NH3]2+, [M+H-H2O]+, [M+H-H2O]+, [M+2H-H2O]2+, [M+3H]3+, [M+CH3COONa+H]+, [M+CH3COONa+Na]+, [M+CH3COONa+NH4]+ | [M-H]-, [M-2H]2-, [M-2H+Na]-, [M-H+Cl]2-, [M-2H+K]-, [M+Cl]-, [2M-H]-, [2M-2H+Na]-, [2M-H+Cl]2-, [2M-2H+K]-, [2M+Cl]-, [M-H-H2O]- |
| dmz | 0.005 | 0.005 | 0.005 |
| main adducts positive | [M+H]+, [M+2H]2+, [M+Na]+, [M+NH4]+ | [M+H]+, [M+2H]2+, [M+Na]+, [M+NH4]+ | [M-H]-, [M+Cl]-, [2M-H]- |
| max charge | 3 | 3 | 3 |
| max isotopes | 4 | 4 | 4 |
| min filter | 2 | 2 | 2 |
| num files | 100 | 100 | 100 |
| polarity | positive | positive | negative |
| ppm | 5 | 5 | 5 |
| files used | 12 | 12 | 12 |
| need optimization | TRUE | TRUE | TRUE |
| noise threshold | 1000 | 1000 | 1000 |
| num iterations | 5 | 5 | 5 |
| number of points | 30 | 30 | 50 |
| ms1 | gap-filled data matrix | gap-filled data matrix | gap-filled data matrix |
| ms2 | fused mgf | fused mgf | fused mgf |
| algorithm | ADAP | ADAP | ADAP |
| noise level ms1 | 1000 | 1000 | 1000 |
| noise level ms2 | 1000 | 1000 | 1000 |
| SN* | 4.57 | 17.38 | 15.6 |
| coefficient area threshold* | 57.94 | 108.56 | 39.05 |
| ms2 mz tol | 0.005 | 0.005 | 0.005 |
| ms2 rt tol | 0.1 | 0.1 | 0.1 |
| noise level | 10000 | 10000 | 10000 |
| peak width (min/max)* | 0.018 / 0.718 | 0.030 / 0.434 | 0.083 / 0.581 |
| rt wavelet (min/max)* | 0.001 / 0.081 | 0.002/ 0.110 | 0.012 / 0.103 |
| peaktable filter | absolute intensity top 30000 | absolute intensity top 30000 | absolute intensity top 30000 |
| dmz* | 0.005 | 0.004 | 0.002 |
| min scan* | 6 | 4 | 4 |
| ppm* | 6 | 12 | 6 |

*value determine via SLAW optimization

**Supplemental Table 4.** Pre-processing steps, input parameters, and set values used for LC-MS data are listed in the order of execution
